# A Coalescent Model of a Sweep from a Uniquely Derived Standing Variant

**DOI:** 10.1101/019612

**Authors:** Jeremy J. Berg, Graham Coop

## Abstract

The use of genetic polymorphism data to understand the dynamics of adaptation and identify the loci that are involved has become a major pursuit of modern evolutionary genetics. In addition to the classical “hard sweep” hitchhiking model, recent research has drawn attention to the fact that the dynamics of adaptation can play out in a variety of different ways, and that the specific signatures left behind in population genetic data may depend somewhat strongly on these dynamics. One particular model for which a large number of empirical examples are already known is that in which a single derived mutation arises and drifts to some low frequency before an environmental change causes the allele to become beneficial and sweeps to fixation. Here, we pursue an analytical investigation of this model, bolstered and extended via simulation study. We use coalescent theory to develop an analytical approximation for the effect of a sweep from standing variation on the genealogy at the locus of the selected allele and sites tightly linked to it. We show that the distribution of haplotypes that the selected allele is present on at the time of the environmental change can be approximated by considering recombinant haplotypes as alleles in the infinite alleles model. We show that this approximation can be leveraged to make accurate predictions regarding patterns of genetic polymorphism following such a sweep. We then use simulations to highlight which sources of haplotypic information are likely to be most useful in distinguishing this model from neutrality, as well as from other sweep models, such as the classic hard sweep, and multiple mutation soft sweeps. We find that in general, adaptation from a uniquely derived standing variant will be difficult to detect on the basis of genetic polymorphism data alone, and when it can be detected, it will be difficult to distinguish from other varieties of selective sweeps.

## Introduction

In recent decades, an understanding of how positive directional selection and the associated hitchhiking effect influence patterns of genetic variation has become a valuable tool for evolutionary geneticists. The reductions in genetic diversity and long extended haplotypes that are characteristic of a recent selective sweep can allow for both the identification of individual genes that have contributed to recent adaptation within a population (i.e. hitchhiking mapping), and for understanding the rate and dynamics of adaptation at a genome-wide level (WIEHE And STEPHAN, 1993; ANDOLFATTO, 2007; EYRE-WALKER And KEIGHTLEY, 2009; ELYASHIV *et al.*, 2014).

While the contribution of many different modes to the adaptive process has long been recognized, early work on the hitchhiking effect focused largely on the scenario where a single co-dominant mutation arose and was immediately beneficial, rapidly sweeping to fixation (MAYNARD SMITH and HAIGH, 1974; Kaplan *et al.*, 1989). Both simulation studies and analytical explorations during the last decade however have drawn attention to models in which adaptation proceeds from alleles present in the standing variation or arising via recurrent mutation once the sweep has already begun (INNAN And KIM, 2004; PRZEWORSKI *et al.*, 2005; HERMISSON and PENNINGS, 2005; PENNINGS and HERMISSON, 2006a,b; Hermisson and PFAFFELHUBER, 2008; BARRETT And SCHLUTER, 2008; RALPH And COOP, 2010; POKALYUK, 2012; ROESTI *et al.*, 2014; WILSON *et al.*, 2014). Collectively, these phenomena have come to be known as “soft sweeps”, a term originally coined by HERMISSON and PENNINGS (2005), and now often used as a catchall phrase to refer to any sweep for which the most recent common ancestor at the locus of the beneficial allele(s) predates the onset of positive selection (MESSER and PETROV, 2013).

Empirical work occurring largely in parallel with the theory discussed above suggests that soft sweeps of one variety or another likely make a substantial contribution to adaptation. For example, many freshwater stickleback populations have independently lost the bony plating of their marine ancestors due to repeated selection on an ancient standing variant at the *Eda* gene (COLOSIMO, 2005), and a substantial fraction of the increased apical dominance in maize relative to teosinte can be traced to a standing variant which predates domestication by at least 10,000 years (STUDER *et al.*, 2011). Additional examples of adaptation from standing variation have been documented in *Drosophila* (MAGWIRE *et al.*, 2011), *Peromyscus* (DOMINGUES *et al.*, 2012) and humans (PETER *et al.*, 2012), among others. Adaptations involving simultaneous selection on multiple alleles of independent origin at the same locus have also been documented across a wide array of species (MENOZZI *et al.*, 2004; NAIR *et al.*, 2006; KARASOV *et al.*, 2010; SALGUEIRO *et al.*, 2010; SCHMIDT *et al.*, 2010; JONES *et al.*, 2013). Nonetheless, the general importance of soft sweeps for the adaptive process remains somewhat contentious (see e.g. JENSEN, 2014; SCHRIDER *et al.*, 2015).

While models of the hitchhiking effect under soft sweeps involving multiple independent mutations have received a fair amount of analytical attention (PENNINGS and HERMISSON, 2006a,b; HERMISSON and PFAFFELHUBER, 2008; POKALYUK, 2012; WILSON *et al.*, 2014), the model of a uniquely derived mutation which segregates as a standing variant before sweeping in response to an environmental change is less well characterized. Present understanding of the hitchhiking effect in a single population under this model comes primarily from two sources. The first is a pair of simulation studies (INNAN And KIM, 2004; PRZEWORSKI *et al.*, 2005), which focused largely on simple summaries of diversity and the allele frequency spectrum, and the second is the general verbal intuition that, similar to the multiple mutation case, the beneficial allele should be found on “multiple haplotypes”. In contrast to the multiple mutation case, these additional haplotypes are created as a result of recombination events during the period before the sweep when the allele was present in the standing variation, rather than due to recurrent mutations on different ancestral haplotypes (BARRETT And SCHLUTER, 2008; MESSER and PETROV, 2013).

Before we turn to the coalescent for sweeps from uniquely derived standing variation, it is worth first asking under what circumstances we might expect such sweeps. To illustrate this we consider a single locus model in which a population that was previously at mutation-drift equilibrium adapts in response to an environmental change, either by drawing on material from the standing variation, or from new mutations which occur after the environmental change. In particular, we are interested in exploring the relationship between the source of genetic material the population uses to adapt and the specific signature left behind in genetic polymorphism data at the conclusion of the event. If adaptation proceeds entirely from *de novo* mutation, the signature will either be that of a classic hard sweep, or a multiple mutation soft sweep. However, if the population adapts at least partially from standing variation, a broad range of possible signatures are possible. First, the population may use more than one allele present in the standing variation, in which case we again have a multiple mutation soft sweep. Alternately, if only a single allele from the standing variation is used, a range of signatures are possible. If the allele was at a frequency less than 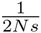 at the moment of the environmental change, then a hard sweep signature is produced because, conditional on escaping loss due to drift and eventually reaching fixation, the allele must have rapidly increased in frequency even before it became beneficial. If adaptation proceeds via an allele that was at some low frequency greater than 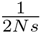, an altered signature is produced (e.g. PRZEWORSKI *et al.*, 2005, and a model for generating that pattern is the primary focus of this paper), whereas adaptation from a single high frequency derived allele leaves essentially no detectable signature in polymorphism data. Drawing on results from a number of previously published studies (HERMISSON and PENNINGS, 2005; PRZEWORSKI *et al.*, 2005; PENNINGS and HERMISSON, 2006a) in the Appendix we calculate the probability of observing each of these different signatures for this model of a sharp transition from drift-mutation equilibrium to positive selection as a function of the population size, strength of selection, and time since the environmental change, and present the results in Figure 1. These calculations reveal that under this model, all of these signatures potentially occur with some probability, and in particular suggests that a sweep of a uniquely derived allele from the standing variation should constitute a non-negligible proportion of all sweeps which begin from mutation-drift equilibrium. Our model also applies to sweeps of alleles which previously exhibited long term asymmetric balancing selection, which represent an unknown fraction of adaptive alleles.

**Figure 1:**
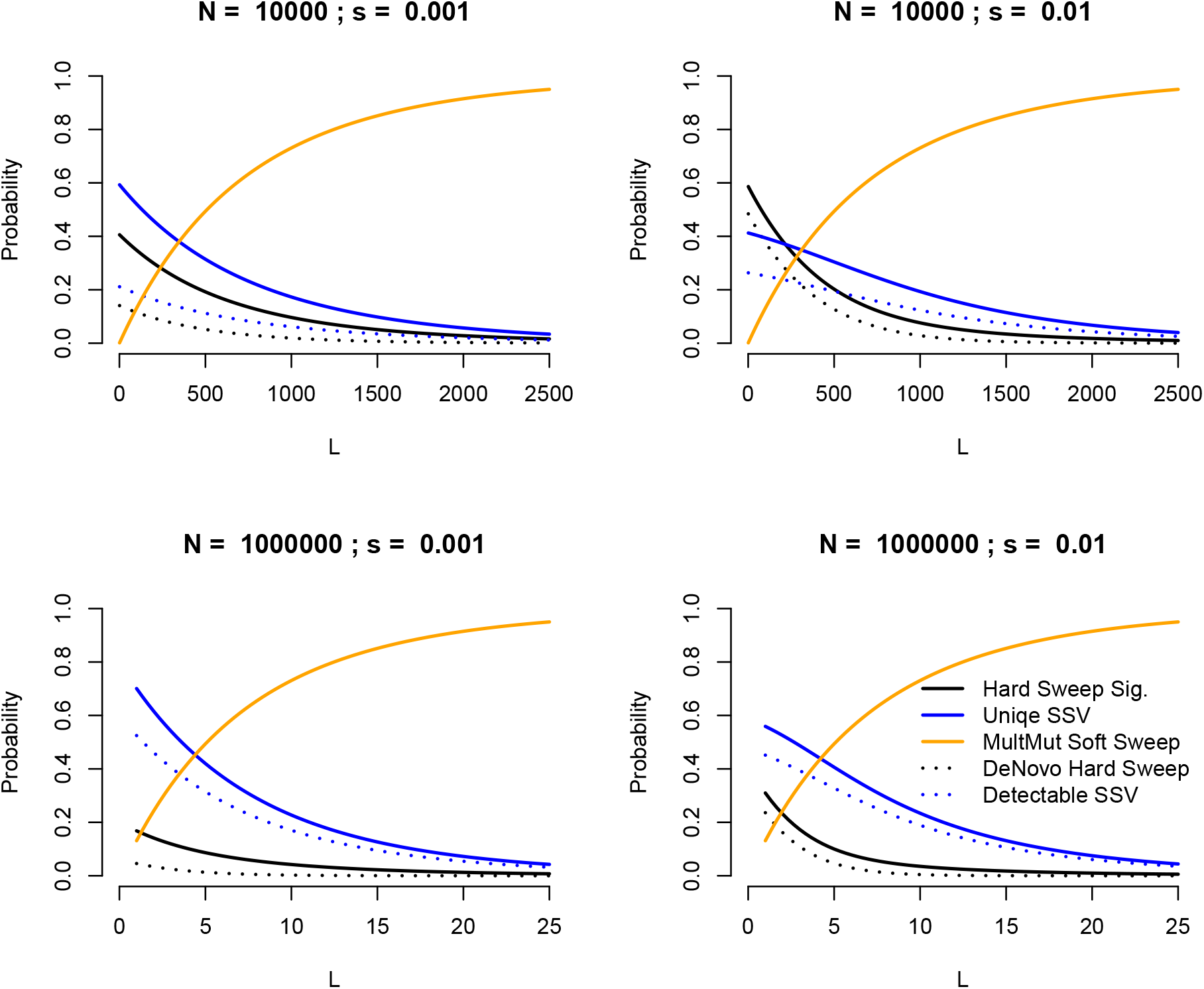
The probability of observing a number of different sweep signatures in a sample of 20 chromosomes assuming a model in which an allele which was previously neutral suddenly becomes beneficial in response to an environmental change. Calculations given in the Appendix. Results displayed for a range of population size (N), selection coefficient (s), mutational target size (L), and assuming 1000 generations since the environmental change. In general, we see that selective sweeps in which adaptation proceeds from a uniquely derived allele represent a non-trivial proportion of all sweeps under this model provided that the mutational target size is not large, and that Ns is not too small. A hard sweep signature is left by any sweep for which a single allele sweeps from a frequency of less than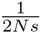, while a Unique Sweep from Standing Variation (SSV) corresponds to any sweep in which a single allele sweeps from a frequency greater than this value. Mutliple mutation soft sweeps refer to the variety described in Pennings and HERMISSON (2006a) and PENNINGS and HERMISSON (2006b). DeNovo Hard Sweep refers sweeps in which the beneficial allele did not arise until after the environmental change (corresponding to the model originally studied by MAYNARD SMITH and HAIGH, 1974), while Detectable SSVs are sweeps of a single unique allele which was present at a frequency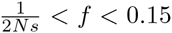, and may therefore plausibly be distinguished from both the hard sweep model and the neutral model.

In this paper, we present an analytical treatment of the model in which an allele with a single mutational origin segregates at a low frequency greater than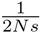(either neutrally or under the influence of balancing selection with asymmetric heterozygote advantage) and then sweeps to fixation after a change in the environment. The central observation is that, with some simplifying assumptions, the recombination events which are responsible for the multiple haplotypes on which the beneficial allele is found have a close analogy to mutations in the infinite alleles model, and we can therefore leverage the EWENS Sampling Formula to obtain an analytical description for the genealogical history of a neutral locus linked to the beneficial allele. We then show that this model can be used to obtain a highly accurate approximation for the expected deviation in the frequency spectrum at a given genetic distance, as well as to shed light on how the expected pattern of haplotype structure differ between the multiple recurrent mutation and sweep from standing variation cases. We conclude with a brief simulation study examining the order statistics of the haplotype frequency spectrum under the classic hard sweep, multiple mutation soft sweep, and standing sweep models, with the aim of demonstrating how future methods to identify and classify sweeps can best make use of this information.

## Model

We consider two linked loci separated on the chromosome by a recombination distance *r*. At one of these loci a new allele, **B**, arises in a background of ancestral **b** alleles. This allele segregates at low frequency for some period of time (either due to neutral fluctuations, or because it is a balanced polymorphism), before a change in the environment causes it to become beneficial and sweep to fixation. A schematic depiction of the model is given in Figure 2. Our aim is to describe some features of genealogies both at the locus of the **B** allele and nearby linked sites, and to use this understanding to build intuition regarding the process of a sweep from standing variation, as well as to derive the patterns of DNA sequence variation we expect to observe near a recently completed sweep from standing variation.

**Figure 2:**
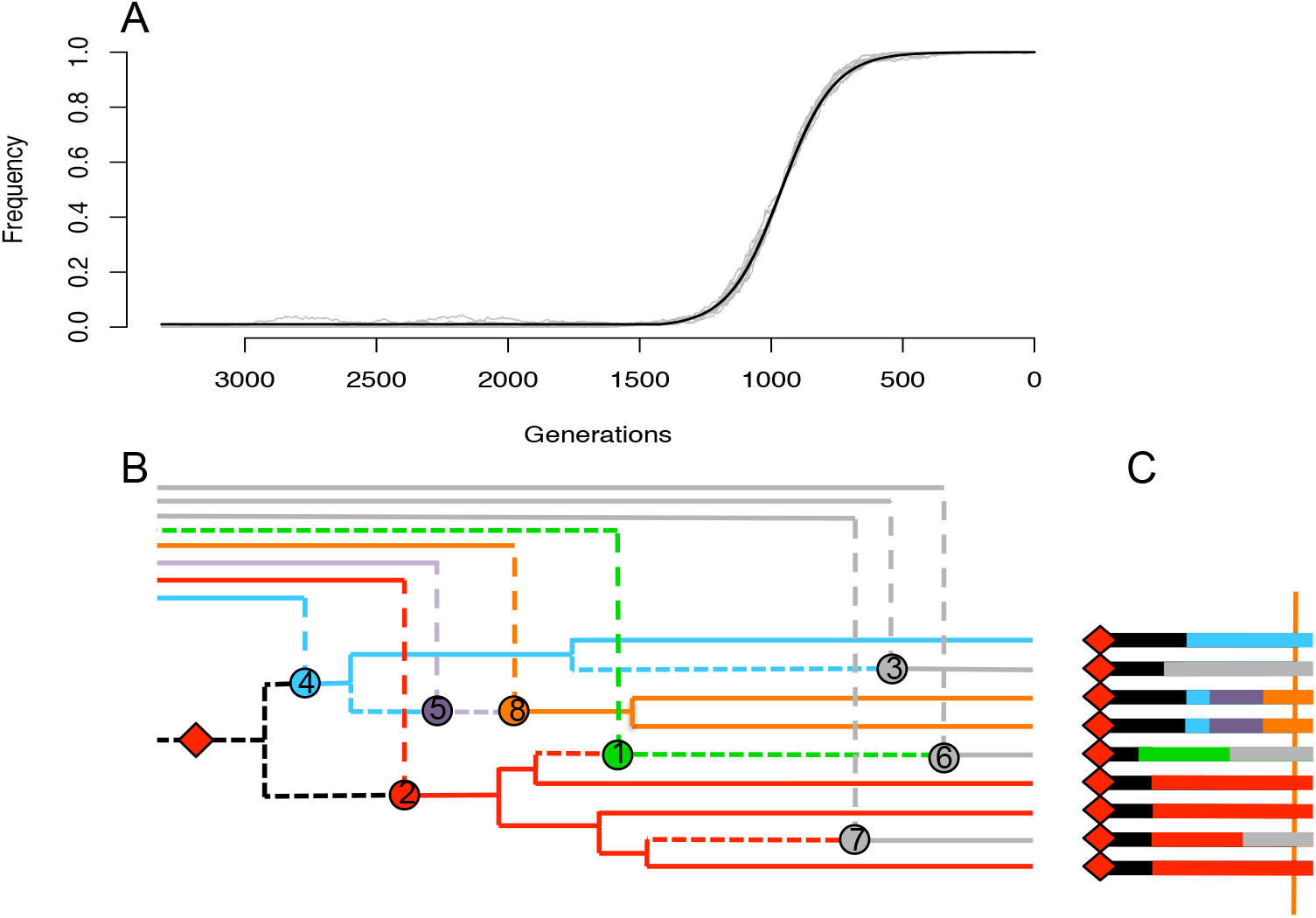
A schematic depiction of our model. (A) Grey lines represent 10 simulated sweeps with *s* = 0.01 and *f* = 0.01 in a population of *N* = 10000. The solid black line represents the frequency trajectory assumed for our analytical calculations. (B and C) The genealogy, history of recombination events, and sequence associated with a sample of nine chromomsomes taken at the moment of fixation. The red dot (on both the genealogy and the sequence) represents the mutation responsible for the beneficial allele. The tree subtending this mutation in panel B is the genealogy at the locus of this mutation. Solid lines represent the genealogy experienced by a neutral site located at the position of the vertical orange bar in panel C, with lineages that escape coalescence under the red mutation coalescing on a longer timescale off the left side of the figure. Stars on the genealogy in panel B represent the recombination events falling between the beneficial mutation and the orange bar in panel C, and are responsible for changes in along the sequence. Short dashed lines represent components of genealogies in between the red mutation and the orange bar which are not experienced at the position of the orange bar, while long dashed lines represent movement from the selected to the non-selected background via recombination. At the distance marked by the orange bar, there are three sweep phase recombinants, and the remaining six sequences are partitioned into three haplotypes of frequencies three, two and one, according to the infinite alleles process described in the main text.

Our general approach is to break the history of the standing sweep into two periods, the first being the time during which the **B** allele is selectively favored and rising in frequency (we refer to this as the sweep phase), and the second being the period after the mutation has arisen but before the environmental shift causes it to become beneficial (we refer to this as the standing phase). We assume that the frequency trajectory of the allele is logistic during the sweep phase, and that selection is sufficiently strong relative to the sample size such that only recombination (i.e. no coalescence) occurs during this phase. We approximate the standing phase by assuming that the frequency of the **B** allele is held at some constant value *f* infinitely far into the past prior to the onset of selection. While this is obviously a coarse approximation to the true history of a low frequency allele, it is nonetheless accurate enough for our purposes, and enjoys some theoretical justification, as we discuss below. The key advantage to using this approximation is that it allows us to model the genealogy of the **B** alleles as a standard neutral coalescent (rescaled by a factor *f*), and therefore to treat recombination events moving away from the selected locus in a manor analogous to mutations in the standard infinite alleles model. This allows us to use a version of the Ewens Sampling Formula to calculate a number of summaries of sequence diversity, and to build intuition for how patterns of haplotype diversity should change in regions surrounding a standing sweep.

## Analysis and Results

### Sweep Phase

Looking backward in time, let *X* (*t*) be the frequency of the **B** allele at time *t* in the past, where *t* = 0 is the moment of fixation (i.e. *X* (0) = 1; *X* (*t*) <1∀ *t* >0). If we consider a neutral locus a genetic distance *r* away from the beneficial allele, the probability that it fails to recombine off of the selected background in generation *t*, given that it has not done so already is 1 − *r* (1 − *X*(*t*)). If we let τ_f_ be the generation in which the environmental change occurred, marking the boundary between the sweep phase and the standing phase (i.e. *X* (τ_*f*_) = *f*), then the probability that a single lineage fails to recombine off the selected background at any point during the course of the sweep phase is given by

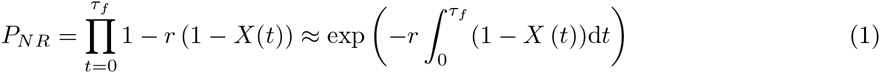

for *r* ≪1. If the effect of our beneficial allele on relative fitness is strictly additive, such that heterozygotes enjoy a selective advantage of *s* and homozygotes an advantage of 2*s*, then the trajectory of the beneficial allele through the population can be approximated deterministically by the logistic function, and the integral in the exponential in equation (1) can be approximated as *ln 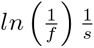*, yielding

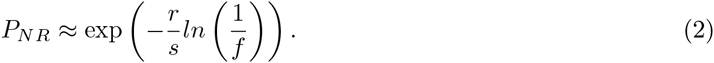

We assume selection is strong, such that there is not enough time for a significant amount of coalescence during the sweep phase. Therefore, each lineage either recombines off the beneficial background, or fails to do so, independently of all other lineages. The probability that *i* out of *n* lineages fail to escape off the sweeping background is then

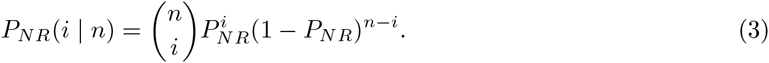

This binomial approximation has been made by a number of authors in the context of hard sweeps (e.g. MAYNARD SMITH and HAIGH, 1974; FAY And WU, 2000; MCVEAN, 2006), but better approximations do exist (BARTON, 1998; DURRETT And SCHWEINSBERG, 2004, 2005; SCHWEINSBERG And DURRETT, 2005; ETHERIDGE *et al.*, 2006; MESSER and NEHER, 2012). Under the hard sweep model, most of the error of the binomial approximation arises due to coalescent events during the earliest phase of the sweep. Because this phase is replaced in our model by the standing phase described below, the binomial approximation is a better fit for our use than in the classic hard sweep case.

### Standing Phase

Looking backward in time, having originally sampled *n* lineages at *t* = 0, we arrive at the beginning of the standing phase at time τ_*f*_ with *i* lineages still linked to the beneficial background, the other *n* − *i* having recombined into the non-beneficial background during the sweep.

We apply a separation of timescales argument, noting that coalescence of the *i* lineages which fail to recombine off the **B** background during the sweep will occur much faster than coalescence of the *n* − *i* lineages which do recombine during the sweep. We therefore assume that nothing happens to lineages on the **b** background until all lineages have escaped the **B** background via either mutation or recombination, at which point **b** lineages follow the standard neutral coalescent.

#### The Coalescent Process of the B Alleles

A number of previous studies have examined the behavior of this process (RANNALA, 1997; GRIFFITHS And TAVARE, 1998, 1999; WIUF And DONNELLY, 1999; WIUF, 2000; GRIFFITHS, 2003; PATTERSON, 2005), either conditional on the frequency of the allele in a sample or in the population. WIUF (2000) has shown that the expected time to the first coalescent event is 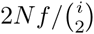 in the absence of other information, e.g. as to whether the allele is ancestral or derived. However, the distribution of coalescence times is no longer exponential. The variance of the time between coalescent events is increased relative to the exponential as a direct result of the fact that the frequency may increase or decrease from *f* before a given coalescent event is reached. Further, in contrast to the standard coalescent, there is non-zero covariance between subsequent coalescent intervals, as a result of the information they contain about how the frequency of the allele has changed, and thus about the rate at which subsequent coalescent events occur. Lastly, if the allele is known to be either derived or ancestral, the expected coalescent times have a more complicated expression, as the allele is in expectation either decreasing or increasing in frequency backward in time due to the conditioning on derived or ancestral status respectively.

Despite these complications, we have found that assuming that all pairs of lineages coalesce at a constant rate 1*/*(2*N f*) and that coalescent time intervals are independent (in other words, that the allele frequency does not drift from *f*) is not a bad approximation when *f* ≪1, even when we condition on the allele being derived (Figures S1, S2 and S3).

The main reason for using this approximation is that, in conjunction with the separation of timescales, it allows us to work with a simple, well understood caricature of the true process (i.e. the neutral coalescent) that still describes the genealogy at the selected site with reasonable accuracy. Given this simplified coalescent process, we can study the recombination events occurring between the beneficial and neutral loci to understand the properties of the genetic variation at the neutral locus that will hitchhike along with the **B** allele once the sweep phase begins.

#### Recombination Events Ocurring During the Standing Phase

We will again rely on the condition that *f* ≪1, and assume that any lineage at the neutral locus that recombines off of the background of our beneficial allele will not recombine back into that background before it is removed by mutation. Under these assumptions, recombination events which move lineages at the neutral locus from the **B** background onto the **b** background can be viewed simply as events on the genealogy at the beneficial locus which occur at rate *r* (1 − *f*) for each lineage independently. Rescaling time by 2*N f*, an understanding of the genealogy at the neutral locus can therefore be found by considering the competing poisson processes of coalescence at rate 1 per pair of lineages, and recombination at rate 2*N rf* (1 − *f*) per lineage.

We are interested in the number and size of different recombinant clades at a given genetic distance from the selected site (colored clades in Figure 2B, which give rise to colored haplotypes in Figure 2C). Under our approximate model for the history of coalescence and recombination at these sites, this a direct analogy of the infinite alleles model (Kimura and Crow, 1964; WATTERSON, 1984). In the normal infinite alleles process, we imagine simulating from the coalescent, scattering mutations down on the genealogy, and then assigning each lineage to be of a type corresponding to the mutation that sits lowest above it in the genealogy. Alternately, we can create a sample from the infinite alleles model by simulating the mutational and coalescent processes simultaneously: coalescing lineages together as we move backward in time, “killing” lineages whenever they first encounter a mutation and assigning all tips sitting below the mutation to be of the same allelic type (GRIFFITHS, 1980).

Given the direct analogy to the infinite alleles model under our set of approximations, the number and frequency of the various recombinant lineage classes at a given distance from the selected site can be found using the Ewens Sampling Formula (EWENS, 1972). The population-scaled mutation rate in the infinitely-many alleles model (*θ/*2 = 2*N µ*), is replaced in our model by the rate of recombination out of the selected class (*R*_*f*_/2 = 2*Nrf* (1 − *f*)). If *i* lineages sampled at the moment of fixation fail to recombine off of the beneficial background during the course of the sweep, then the probability that these *i* lineages coalesce into a set of *k* recombinant lineages is

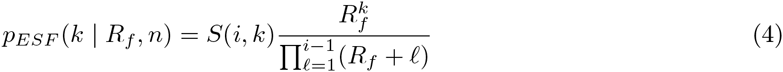

where *S*(*i, k*) is an unsigned Stirling number of the first kind

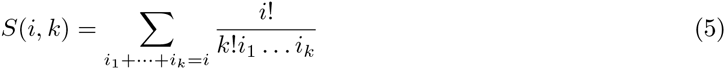

These recombinant lineages partition our sample up between themselves, such that each lineage has some number of descendants in our present day sample 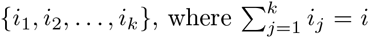 Conditional on *k*, the probability of a given sample configuration is

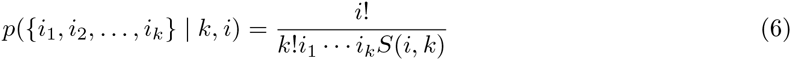

Note that this does not depend on *R*_*f*_, which gives the classic result that the number of alleles is sufficient statistic for *R*_*f*_ (i.e. the partition is not needed to estimate *R*_*f*_). Figure 3 shows a comparison of this approximation to simulations for the number of distinct coalescent families at a given distance from the focal site.

**Figure 3:**
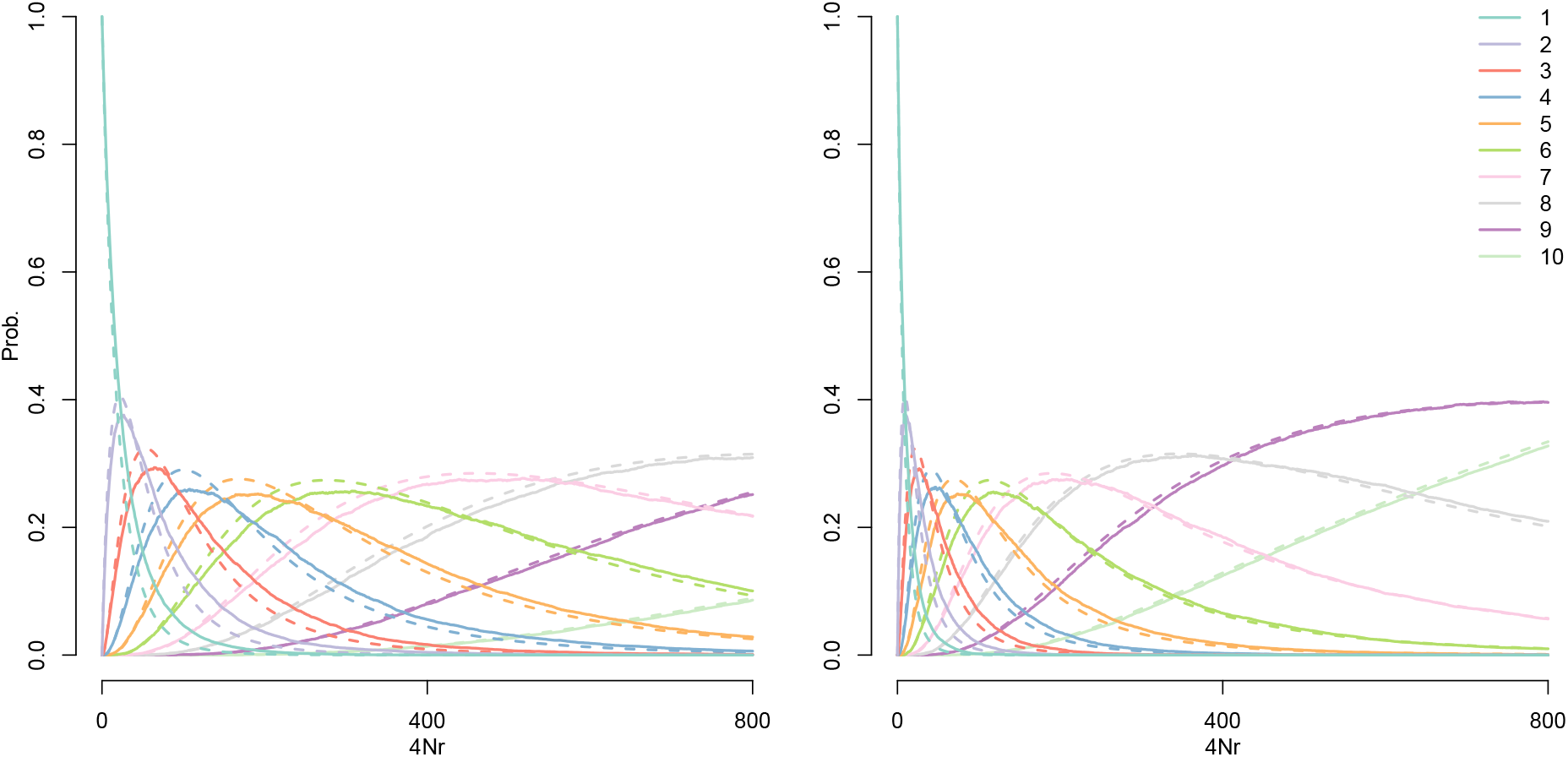
The probability that a sample of 10 lineages taken on the background of an allele at frequency 1% (panel A) or 5% (panel B) coalesce into k families before exiting the background, as a function of population-scaled genetic distance (4*N r*) from the conditioned site. The effective population size in the simulations is *N* = 10000. The solid lines give the proportion of 1000 coalescent simulations, with an explicit stochastic frequency trajectory (as describe in the Simulation Details section), in which *k* families of lineages recombined off of the sweep at distance 4*N r*. The dotted lines give our approximation under the Ewens Sampling Formula (eqn (4)) with *R*_*f*_ = 4*N rf* (1 − *f*).

We will usually be interested in the case where *f* ≪ 1, thus *R*_*f*_ ≈ 4*N f r*. As such, the properties of the standing part of the sweep are well captured by the population sized-scaled compound parameter 4*N f*, the number of individuals who carry the selected allele when the sweeps begins. This means that the effect of standing variation on sweep patterns depends critically on the effective population size. A sweep from a variant at frequency 1*/*5000 would for all intents and purposes be a hard sweep in humans, where the historical effective population size is around 10 thousand, but would result in quite different patterns in *Drosophila melanogaster*, whose long-term effective population size is closer to 1 million.

### Patterns of neutral diversity surrounding standing sweeps

This approximate model of the coalescent for a sweep from standing variation allows us to calculate a number of basic summaries of sequence variation in the region surrounding the sweep. For now we neglect mutations which occur over the time-scale of our shrunken coalescent tree, and assume that all diversity comes from mutations that occurred prior to the sweep, or equivalently that this part of the genealogy contributes negligibly to the total time. This corresponds to an assumption that 2*N* ;Cf ≪ 2*N* ;, in line with our previous assumption that *f* ≪ 1. So long as this assumption holds, we can consider patterns of diversity in our sample at a given site simply by considering properties of the recombinant lineages in our sample, which correspond to alleles drawn independently from a neutral population prior to the start of our sweep. We partially relax this assumption in the appendix for those of our calculations where it substantially affects the fit to simulation data.

#### Reduction in Pairwise Diversity

The expected reduction in pairwise diversity following a standing sweep relative to neutral expectation is given by the probability that at least one lineage in a sample of two manages to recombine off of the **B** background (during either the sweep phase or the standing phase) before the coalescent event during the standing phase

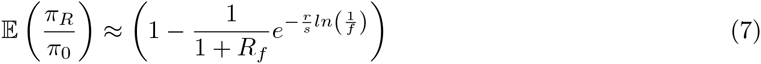

(Figure 4). Given the exponential form of *P*_*NR*_, and the fact that 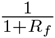 can be approximated as *e*^*-R*^_*f*_for small values of *R*_*f*_, we can further approximate (7) as 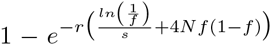. Recalling that the reduction in diversity for a classic hard sweep with strong selection can be approximated as 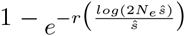 (DURRETT And SCHWEINSBERG, 2004; PENNINGS and HERMISSON, 2006b), it is tempting to suppose that there may exist a choice of Ŝ an “effective” selection coefficient, for which the classic hard sweep model produces a reduction in diversity over the same scale as the standing sweep model. While it is simple to set the the terms in the exponentials equal to one another and solve for the appropriate value of 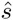 (see Appendix), it turns out that for all choices of *N*_*e*_, *s*, and *f* for which our model applies, 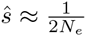. In other words, the reduction in diversity caused by a sweep from standing variation cannot be caused by a hard sweep in which standard strong selection approximations apply, which means that care should be taken when interpreting estimates of the rate of adaptation which depend solely on classic hard sweep strong selection approximations (ELYASHIV *et al.*, 2014). No doubt there is a choice of selection coefficient under which a weakly selected allele will produce a similar reduction in diversity, but there are no adequate approximations available under this model, and we do not pursue it further here.

**Figure 4:**
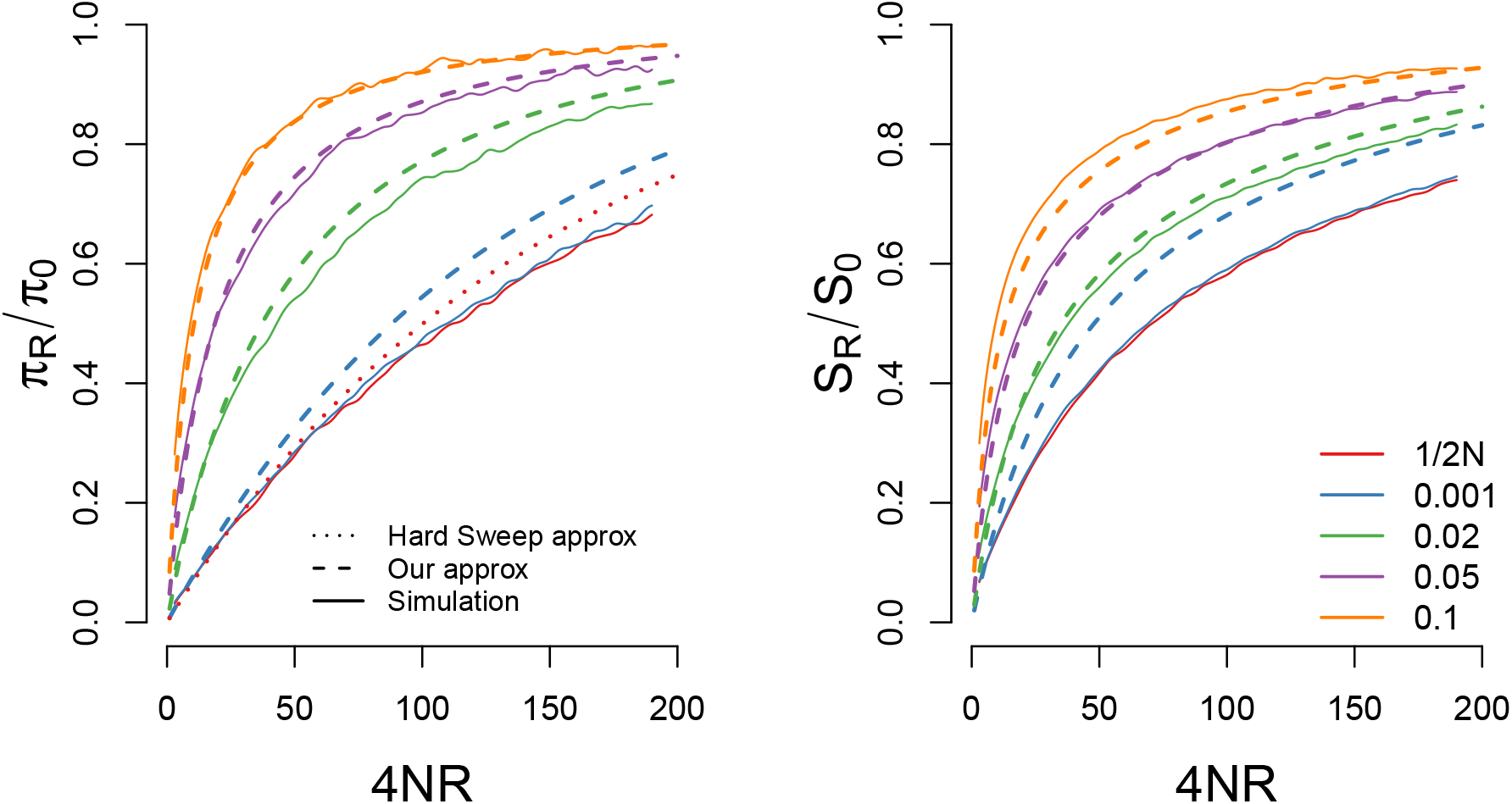
A comparison of our approximations for the reduction in (A) pairwise diveristy and (B) the number of segregating sites for a sweep with *s* = 0.05 and *N* = 10000 starting from a variety of different frequencies. For pairwise diversity we also include the hard sweep approximation given in eqn (26). Our approximations are generally accurate so long as the sweep begins from a frequency greater than 1*/*2*N s*

#### Number of Segregating Sites

We can also use our approximation to calculate the total time in the genealogy at a given distance from the selected site, which allows us to calculate the expected number of segregating sites. Conditional on *m* independent lineages escaping the sweep, the expected total time in the genealogy is 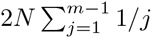, the standard result for a neutral coalescent with *m* lineages (WATTERSON, 1975). For a moment conditioning on no recombination during the sweep phase, the probability that *k* independent lineages escape during the standing phase, given that there are at least 2 (otherwise all coalescence occurs on the **B** background and *T*_*T*__*OT*_ ≈ 0) is

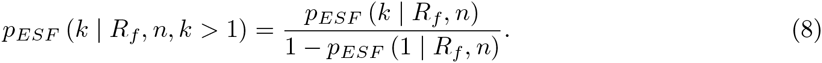

Conditional on no recombination during the sweep phase, the expected time in the genealogy is

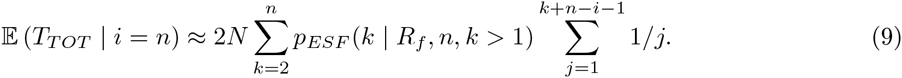

When we allow for all possible numbers of singleton recombinants during the sweep phase, the expected total time in the genealogy is

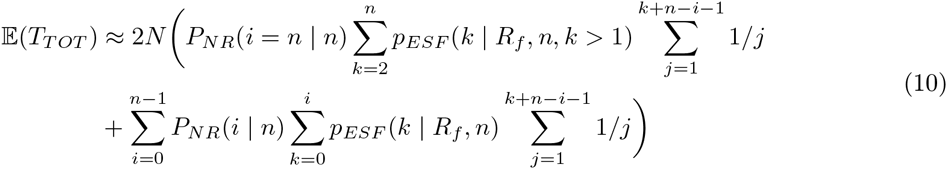

(note that we have taken *p*_*ESF*_ (0 *R*_*f*_, 0) = 1, and *p*_*ESF*_ (0 *R*_*f*_*, i*) = 0 ∀ *i >* 0; whereas it is typically impossible to obtain a sample with zero alleles, in our case we must define these probabilities to accommodate the case in which all *n* lineages recombine out during the sweep phase) and the expected number of segregating sites can be found by multiplying this quantity by the mutation rate (Figure 4).

#### The Frequency Spectrum

Finally, we can use our approximation to obtain an expression for the full frequency spectrum at sites surrounding a sweep from standing variation. To break the problem into approachable components, we first consider the frequency spectrum of an allele that is polymorphic within the set of lineages which do not recombine during the sweep (ignoring it ' s frequency in the sweep phase recombinants), and we condition on a fixed number *k* recombinant families from the standing phase. Borrowing from PENNINGS and HERMISSON (2006b) (equation 14 of their paper), if we condition on *j* out of these *k* recombinant lineages carrying a derived allele, then we can obtain the probability that *l* of the *i* sampled lineages carry the derived allele by summing over all possible partitions of the *i* lineages into *k* families such that the *j* recombinant ancestors carrying the derived mutation have exactly *l* descendants in the present day

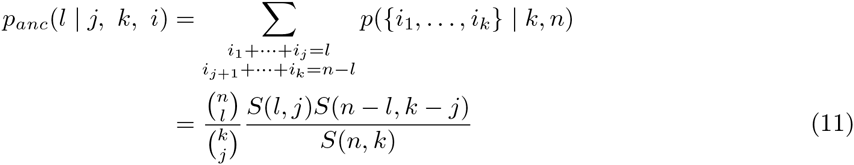

Next, we write *q* (*j* |*k*) to denote the number of polymorphic mutations that were present *j* times among the *k* ancestral lineages which escape the standing phase. For our purposes, we will assume this follows the standard neutral coalescent expectation

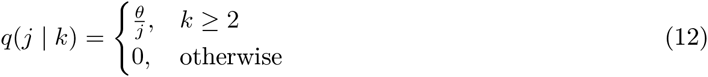

although an empirical frequency spectrum measured from genome-wide data, as in Nielsen *et al.* (2005), could also be used. The expected number of derived alleles that are present in *l* out of *i* sampled lineages, conditional on there having been *k* recombinant families, is then

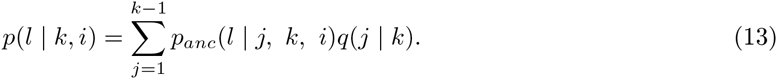

Summing over the distribution of *k* given by (4), we obtain an expression for the frequency spectrum within the set of *i* lineages which do not recombine during the sweep as

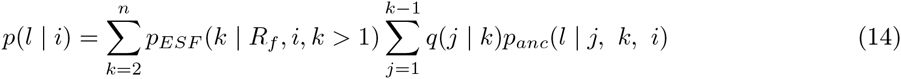

This expression is essentially identical to the one presented in equation 15 of PENNINGS and HERMISSON (2006b). The only difference is that the Ewens clustering parameter in their model is given by the beneficial mutation rate, and holds only for sites fully linked to the selected loci, whereas in our model it is a linear function of the genetic distance from the selected site. In terms of accurately describing observed patterns of polymorphism, this approximation is highly accurate for loci that are distant from the focal site, but breaks down for loci that are tightly linked. The reason for this is that very near the focal site, it is actually very unlikely that there have been any recombination events at all, and so while polymorphism is rare, when it is present it is likely to have arisen due to new mutations on the genealogy of the **B** allele (in which case their distribution is that of the standard neutral frequency spectrum), rather than ancestrally. While a full accounting for the contribution of all new mutations under this model is beyond our scope, we can develop an *ad hoc* approximation by assuming that mutations are new if there have not been any recombination events, and are old if there has been at least one recombination (see Appendix). This approximation is quite accurate, especially when the focal allele is at low frequency (Figure 5).

When we allow for recombination during the sweep, the expression becomes more complex, as we must take into account the fact that a mutation may be polymorphic after the sweep even if it is either absent or fixed in the set of lineages which hitchhike. Nonetheless, we obtain an expression for the frequency spectrum of ancestral polymorphism as

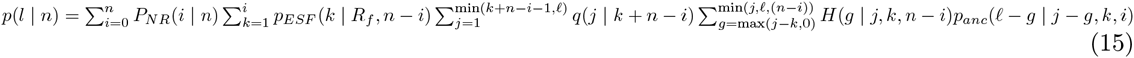

where

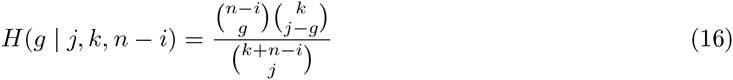

gives the probability that *g* out of a total of *j* derived alleles which existed before the sweep are found on singleton recombinants created during the sweep, given that there are *n* − *i* singletons, and *k* recombinant families created during the standing phase.

In words, *n* − *i* lineages recombine out during the selected phase, while the remaining *i* lineages are partitioned into *k* families at frequencies {*i*_1_,…, *i*_*k*}_ due to recombination and coalescence in the standing phase. Out of the *n* − *i* singleton lineages, *g* of them carry the derived allele, while the remaining *j − g* copies of the derived allele give rise to *l − g* derived alleles due to coalescence during the standing phase, and we take the sum over all possible combinations of these values which result in a final frequency of 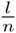in the present day sample.

Once again, this expression is accurate far from the selected site, but in error at closely linked sites due to the contribution of new mutations. Again, we can develop an *ad hoc* approximation by allowing for new mutations during the sweep phase on any lineage that does not recombine during that phase, and on the *i* lineages that reach the standing phase, provided there are no recombination events during that phase (see Appendix and Figure 5). New mutations during the standing phase are ignored once there has been at least one recombination event during that phase. This approximation is quite accurate at all distances (especially when the sweep comes from relatively low frequency), and highlights the fact that sweeps from standing variation are characterized by an excess of derived mutations at a range of frequencies greater that 40-50%, in contrast to hard sweeps, which exhibit a much stronger skew towards extremely low or high frequency alleles (PRZEWORSKI *et al.*, 2005).

**Figure 5:**
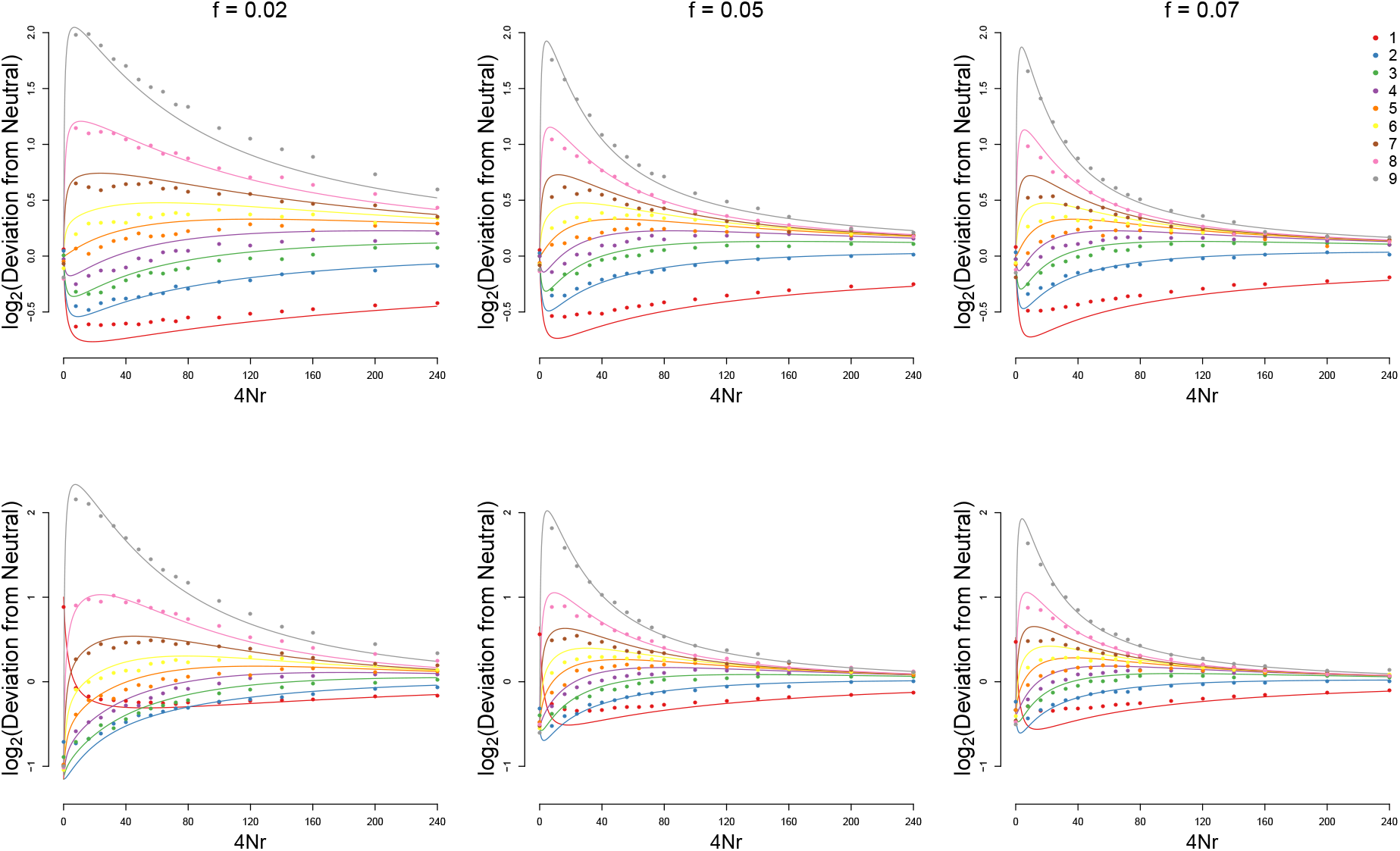
The frequency spectrum, in a sample of n = 10 in a population of N = 10000, for a neutral allele sampled on the background of the beneficial allele either immediately before it sweeps (A,B,C) or immediately after fixation (D,E,F). Results are shown as the log ratio of the normalized frequency relative to its expectation under the standard neutral coalescent. s = 0.05 for the post fixation case.

#### Patterns of Haplotype Variation and Routes to Inference

To this point, we have focused on an analytical description for the effect of a sweep from standing variation on a single tightly linked neutral locus on the same chromosome. It is also of value to consider the effect of a sweep as a process that occurs along the sequence, as this gives some perspective into how haplotype structure unfolds in the region surrounding a standing sweep. Efforts to identify standing sweeps via polymorphism data hinge on identifying recombination events which occurred during the standing phase by recognizing the way in which they break a single core haplotype down into a succession of coupled samples from the Ewens’ distribution with progressively larger clustering parameters. We first describe some properties of pairs of sequences, before considering larger samples.

For any pair of sequences, recombination events from the sweep phase are encountered at rate *ln 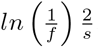*, while events from the standing phase are encountered at rate 2*N f*. A simple measure of the relative importance of the two phases for patterns of haplotype structure and LD can be found by competing Poisson processes. The probability that the first recombination encountered traces its history to the standing phase is

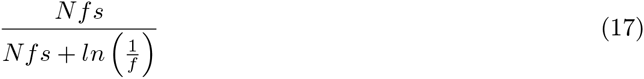

and in general events occurring during the standing phase will dominate the haplotype partition when 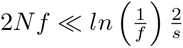 while those from the sweep phase will dominate when 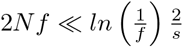.

Next consider that the lower bound on the frequency from which a sweep can start and still conform to our model is 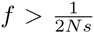. Below this frequency, the effect of conditioning on fixation is so strong that the shape of the genealogy is much more similar to that expected under the classic hard sweep model. However, conditional on a given selection coefficient, most sweeps from standing variation are likely to come from frequencies close to this stochastic threshold, where the probability that the first event from the standing phase occurs before the first event from the sweep phase is

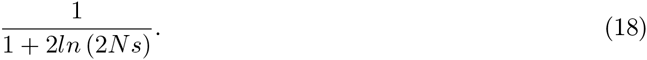

Therefore, while increases in 2*N s* result in an increased probability of sweeps conforming to our model relative to the classic hard sweep model (Figure 1), these sweeps become more and more difficult to distinguish from classic hard sweeps due to the standing phase’s weakened effect when the sweep begins from lower frequencies. While these sweeps from low frequency standing variation can at least in principle be distinguished from classic hard sweeps on the basis of polymorphism data, the task is difficult and requires relatively large sample sizes.

The practical task of identifying a sweep from standing variation requires more extensive knowledge about the haplotype partition from a larger sample. The necessary task is to identify recombination events from the standing phase as they unfold along the sequence (see Figure 2C). Unfortunately, explicit analytical expressions for these haplotype partition transitions are unavailable under any sweep model, and ours is no exception (although INNAN and NORDBORG, 2003, have provided some results regarding our standing phase). Nonetheless, we gain a few simple insights from a description of the process, and this descriptions motivates some further simulations.

If we consider the best case for identifying a sweep from standing variation and distinguishing it from a classic hard sweep, this should occur in the parameter regime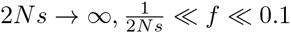, where events from the sweep phase tend to happen much more distantly than those from the standing phase, but the effect of the standing phase still unfolds gradually enough that it can be distinguished from neutrality.

In this limit, the sweep happens instantaneously, and all time in the tree is equal to the total time from the standing phase *T*_*stand*_. The distance to the first recombination is ≈ exp (*r*_*BP*_ (1-*f*) *T*_*stand*_). Using the standard approximation for the total time in the tree, the expected length scale over which a single haplotype should persist away from the selected site is ≈ 1*/*(2*N*_*e*_*r*_*BP*_ *f* (1-*f*) *log*(*n-*1)) (and twice this distance if we consider both sides of the sweep). This recombination partitions the haplotypes according to the standard neutral frequency spectrum (e.g. the green recombinant moving to the left in Figure 2C). Moving down the sequence we then generate the next distance to a recombination, again from ∽ exp (*r*_*BP*_ (1-*f*) *T*_*stand*_). We again uniformly simulate a position on the tree for this new recombination, however, this time only a recombination on some of the branches would result in a new haplotype being introduced into the sample (e.g. the red recombinant in Figure 2B is responsible for the second transition in the haplotype partition scheme in Figure 2C). If the recombination falls in a place that doesn’t alter the configuration we ignore it and simulate another distance from this new position. Otherwise, we keep the recombination event and split the sample configuration again. (For example, the orange recombinant in Figure 2C does not alter the status of identity by descent relationships with respect to the sweep, and therefore does not result in an increase in the number of haplotypes under our convention.) We iterate this procedure moving away from the selected site, generating exponential distances to the next recombination, placing the recombination down, updating the configuration if needed, until we reach the point that every colored haplotype is a singleton. We then repeat this procedure on the other side of the selected site using the same underlying genealogy.

An equivalent way to describe this process is to simulate distances to the next recombination that alters the configuration, given the tree and the previous recombinations. To do this we consider the total time in the tree where a recombination would alter the configuration. Numbering these recombinations out from the selected site, we start at the selected site *i* = 0, with *T*_0_ = *T*_*stand*_ and generate a distance to the first recombination ∽ exp(*r*_*BP*_ (1 − *f*) *T*_0_). We place the recombination on the tree, then prune the tree of branches where no further change in sample configuration could result in a new colored haplotype. We then set *T*_*i*_ to the total time in these pruned subtrees, place the next recombination uniformly on the pruned branches at an ∽ exp(*r*_*BP*_ (1 − *f*) *T*_*i*_) distance and carry on this process till we have pruned the entire tree such that all lineages reach their own unique recombination event before coalescing with any other lineage.

#### Routes to Inference

Any effort to identify and distinguish sweeps from standing variation is necessarily an attempt to identify these characteristic recombination events (and also potentially the new mutations which occur along the genealogy according to essentially the same process). Effectively what we would like is an analytical way to describe how the infinite alleles model “unfolds” along the sequence as we increase the size of the locus at one end. As discussed above, marginally at a given site the partitioning of haplotypes (integrating out the unobserved genealogy) is given by the ESF, but it is unclear to us how to couple together the partitioning at two sites in an analytically tractable way. For example, efficient ways of computing

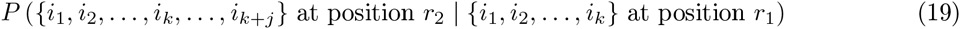

and related expressions, in combination with some of the machinery we use above for our frequency spectrum calculations could in principle be used to take a product of approximate conditional (PAC) likelihoods approach to inference under different sweep models. Of course, such computations could be done via brute force numerical integration over all possible genealogies, but this is unfeasible for any sample size large enough to be useful. One would need to be able to calculate such quantities under multiple different sweep models in order to deploy them, and so this would require further development under all three models. A more direct (and perhaps accurate) route to inference may actually be to build upon recent developments in coalescent HMMs (LI and DURBIN, 2011; PAUL *et al.*, 2011; SHEEHAN *et al.*, 2013; RASMUSSEN *et al.*, 2014) and explicitly model the effect of selective sweeps on the ancestral recombination graph. Our model suggests a way to accomplish this effectively for sweeps from standing variation, and recent work on both hard and soft sweeps (BARTON, 1998; DURRETT and SCHWEINSBERG, 2004, 2005; SCHWEINSBERG And DURRETT, 2005; ETHERIDGE *et al.*, 2006; HERMISSON and PFAFFELHUBER, 2008; MESSER and NEHER, 2012) provide a route to doing so under these models. A third approach, which we explore below, is to continue along the lines of popular sweep finding approaches implemented to date, and define summary statistics which can effectively distinguish between different models. Nonetheless, while both of these approaches seem more likely to be fruitful than the PAC approach described above, expressions like (19) would represent significant progress in relating the infinite alleles and infinite sites models, and given the general interest in the ESF motivated by exchangeable partitions and clustering algorithms, are likely of wider interest.

### Observed haplotype frequency spectum

To this point, our discussions of haplotype variation have focused on haplotypes defined via identity-by-descent, which cannot be observed directly. It is useful to consider how the understanding gained here can be leveraged to improve our ability to identify standing sweeps. To do this we turn to the ordered haplotype frequency spectrum.

For a window of size *L*, we define the the ordered haplotype frequency spectrum (OHFS) as *ℋ*_*L*_ = *{h*_1_*, h*_2_*,…, h*_*H*_*L }*, where *h*_*p*_ gives the sample frequency of the *p*_*th*_ most common haplotype and there are a total of *H*_*L*_ distinct haplotypes within the window. Coarse summaries of the OHFS have been a popular vehicle for sweep-finding methods (e.g. EHH, iHS and H12: SABETI *et al.*, 2002; VOIGHT *et al.*, 2006; GARUD *et al.*, 2015; GARUD and ROSENBERG, 2015). We focus on identifying which aspects of the OHFS should be most informative about the size, as well as the shape of the genealogy at the focal site.

Specifically, we conducted coalescent simulations with a sample size of *n* = 100 chromosomes under four different models of sequence evolution (hard sweeps, standing sweeps from *f* = 0.05, soft sweeps conditional on three origins of the beneficial mutation, and neutral), with all sweep simulations set to *s* = 0.01. For the standing sweeps, this corresponds to 4*N sf* = 20, and thus to a scenario where the standing phase has a slightly stronger, but not overwhelming, impact on the final patterns of haplotype diversity.

One simple prediction on the basis of our analytical investigation above is that, because the genealogy under a standing sweep is generally larger than that under a hard sweep, recombination events out of the sweep should accumulate more quickly along the sequence and therefore the total number of haplotypes in a window of a given size should be larger for a standing sweep than for a hard sweep. In Figure 6 we show from simulations *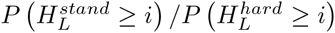* over a range of *L* for one sided windows extending away from the selected site. As expected, we see that the number of haplotypes increases more quickly with distance from the selected site for a standing sweep than for a hard sweep. Unfortunately, as we alluded to above, this signal is largely confounded by the fact that one can also obtain a similarly rapid increase in the number of haplotypes from a hard sweep with a slightly weaker selection coefficient.

**Figure 6:**
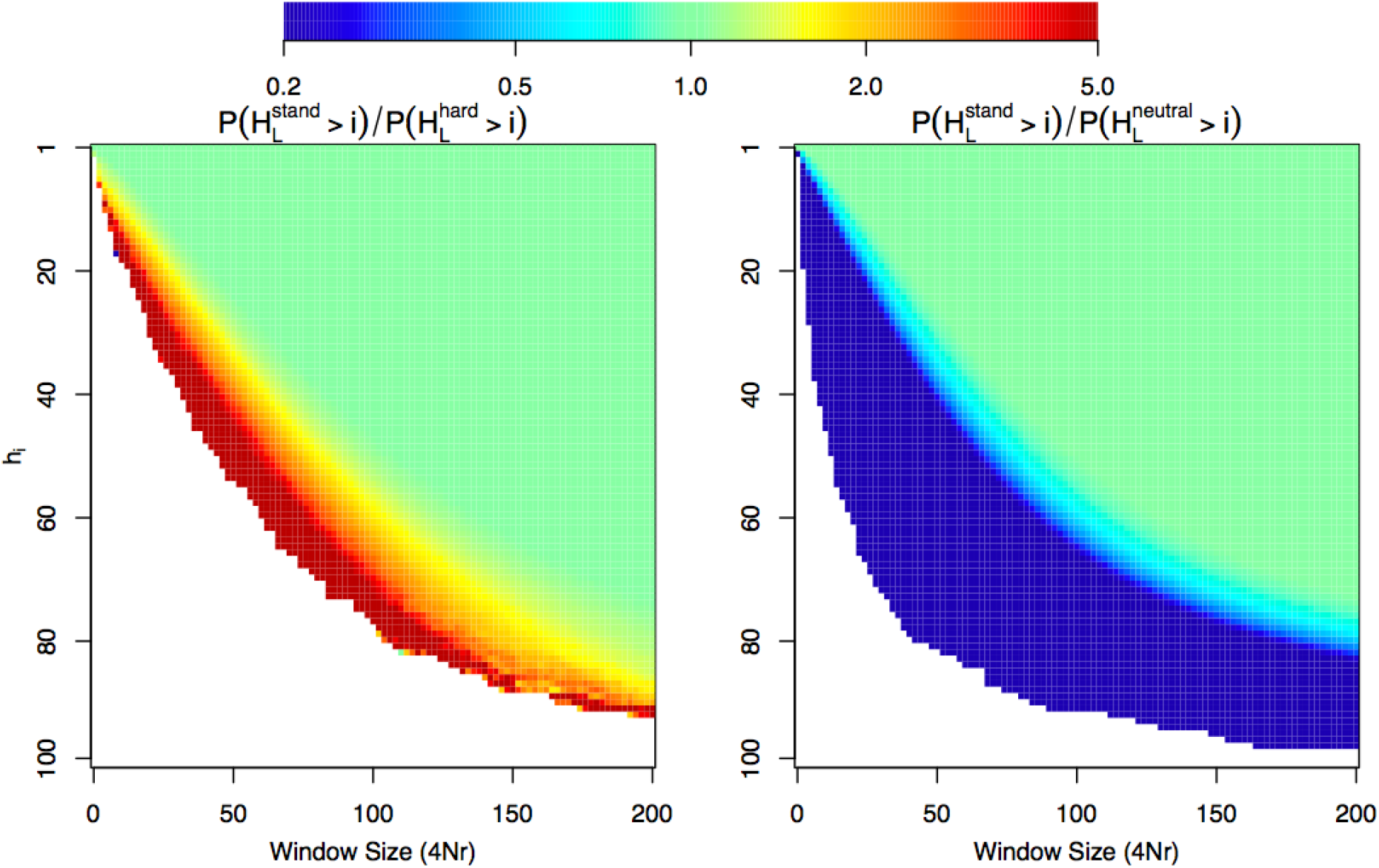
The ratio of the probability that there are at least *i* haplotypes in a one sided window extending away from the selected site for the standing sweep model relative to the hard sweep model (A) and the neutral model (B). For all simulations *n* = 100, *N* = 10000, and we simulate a chromosomal segment with total length 4*N r* = 200 divided into 500, 000 discrete loci, with 4*N μ* = 200 for the whole segment.

Alternately, we may hope to use to OHFS to obtain information about the shape of the genealogy at the focal site, which should hopefully be less confounded by a change in selection coefficient. This information is found in differences in the relative frequencies of certain haplotypes between the different models, rather than in the total number of haplotypes. Specifically, if there are multiple haplotypes present close to the selected site, they should be more common under the sweeps from standing variation model than the full sweep model. In other words, there should be a window near the selected site where 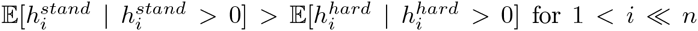, owing to recombinations occurring on internal branches of the genealogy from the standing phase.

In Figure 7, we 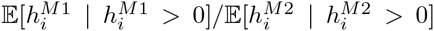 where *M* 1 and *M* 2 denote different models of sequence evolution (i.e. standing, hard, or soft sweep or neutral). In particular, we wish to draw attention to the fact that, similar to the multiple mutation model, most of the useful information within the OHFS for distinguishing standing sweeps from hard sweeps lies within a small window near the selected site, and comes in the form of a decrease in the relative frequency of the core haplotype, and corresponding overabundance of the next few most common haplotypes. In contrast to the multiple mutation case, the enrichment of moderate frequency haplotypes for the standing case is relatively subtle, and beyond moderate recombination distances, there is little information to distinguish a standing sweep from a hard sweep. We also observe that, contrary to multiple mutation soft sweep model, far away from the selected site, standing sweeps resemble hard sweeps across the majority of the OHFS. This is in line with expectations from our results above, in that close to the selected site the haplotype partition is dominated by events occurring during the standing phase, while far from the selected site it is dominated by events occurring during the sweep phase.

**Figure 7:**
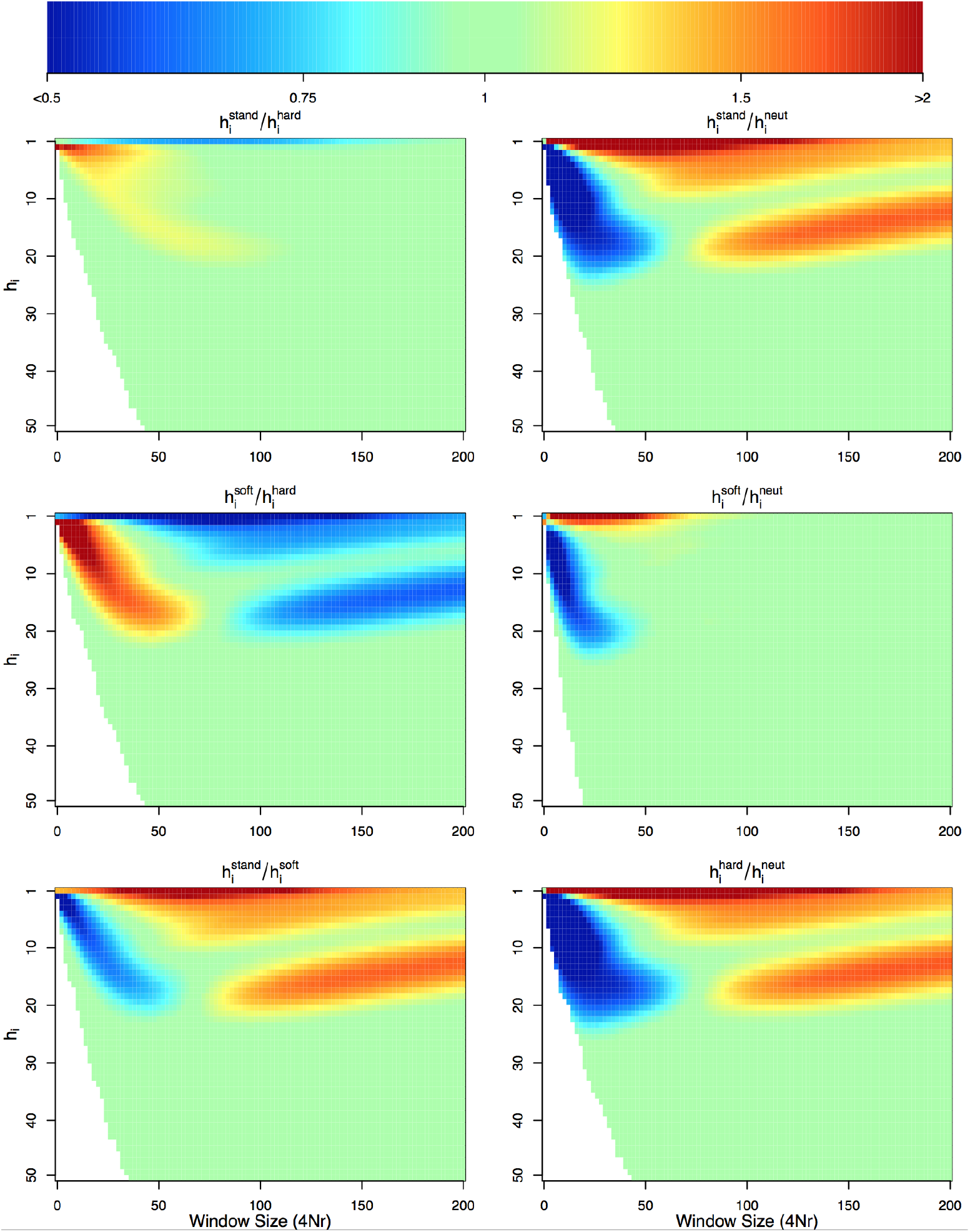
The ratio of expected sample frequency of the *i*^*th*^ most common haplotype, conditional the haplotype existing in our sample, between two different models *M* 1 and *M* 2, i.e.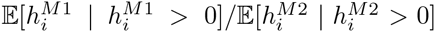. This is shown as a function of the recombination distance from the selected site. Green indicates that the frequency of the *i*^*th*^ most common haplotype is similar in the two models, blue that it has lower frequency under model *M* 1, red that it has lower frequency in *M* 2. We simulated coalescent histories for sample size of *n* = 100 chromosomes under four different models of sequence evolution hard sweeps, standing sweeps from *f* = 0.05, soft sweeps conditional on three origins of the beneficial mutation, and a neutral model, with all sweep simulations using a selection coefficient of *s* = 0.01. For all simulations *N* = 10000, and we simulate a chromosomal segment with total length 4*N r* = 200 divided into 500, 000 discrete loci, with 4*N μ* = 200 for the whole segment.

On the basis of these simulation results, we suspect that future methods for identifying and distinguishing different varieties of sweeps will see benefits from incorporating haplotypic information over a range of window sizes surrounding the focal site, and from pushing deeper into the haplotype frequency spectrum, particularly when large samples are available.

## Discussion

An accurate portrait of the patterns of sequence diversity expected in the presence of recent or ongoing positive selection has proven to be vital for the identification of adaptive loci. Recent theoretical and empirical work has drawn attention to the fact that adaptation from standing variation may be relatively common, and that patterns of sequence variation produced in such scenarios may differ markedly from those produced under the classical hard sweep model. In this paper, we’ve focused on developing a tractable model of strong positive selection on a single mutation which was previously segregating (or balanced) at low frequency. We have shown that many aspects of the pre-sweep standing phase of the mutation’s history on post-hitchhiking patterns variation can be approximated via an application of the Ewens Sampling Formula. This provides a way to build intution for the process and obtain various analytical approximations for patterns of variation following a sweep from standing variation.

Our results can be understood within the context of a number of recent approximations for different sweep phenomena, which divide the sweep into distinct phases (see e.g. BARTON, 1998; ETHERIDGE *et al.*, 2006). Because rates of coalescence and recombination vary across these phases, different sections of the sequence surrounding a sweep will convey information about different phases, with sites distant from the selected locus generally conveying information about the late stages of the sweep, and sites close to the selected locus conveying information about the early stages of the sweep. In our model, the late phase corresponds largely to what we’ve called the sweep phase, while the early phase corresponds to our standing phase. In general, the major differences between different sweep phenomena occur during the earliest phases, and thus the information to distinguish them is found near to the selected site, while extra information about the strength of the sweep can be found at sites that are more distant.

It is worth noting however that all of our results are obtained for populations with equilibrium demographics. If population size is variable, particularly over the course of the standing phase, then the ESF fails to accurately describe recombinations during this phase, just as it fails to accurately describe the infinite alleles model with non-equilibrium demographics. Inference methods based on our analytical calculations would likely be inaccurate in these situations (BANK *et al.*, 2014). Nonetheless, the general insight remains that, holding demography equal, sweeps from standing variation will generate genealogies with longer internal branches than classic hard sweeps, and will therefore be characterized by more intermediate frequency haplotypes.

Although we do not pursue it, essentially all of our results also likely apply to fully recessive sweeps from *de novo* mutation. This is because a recessive beneficial mutation is effectively neutral until it reaches sufficient frequency for homozygotes to be formed at appreciable enough rates to feel the effects of selection. The result is that recessive sweeps should be fairly well approximated by setting 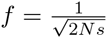 in our model for the standing phase and taking the value of *P*_*NR*_ for the latter phase to be

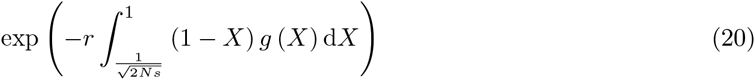

where *g* (*X*) is the Green ' s function for a recessive allele under positive selection. This conclusion is foreseen by EWING *et al.* (2011), who suggested just this sort of approximation for the reduction in diversity following a recessive sweep. The result is that it is likely to be extremely difficult to distinguish between a sweep from previously neutral or balanced standing variation and a recessive sweep without additional biological information about dominance relationships at the locus of interest.

Unfortunately, our work largely confirms the intuition and existing results indicating that standing sweeps are likely to be rather difficult to identify, and characterize, on the basis of genetic data from a single population time-point, and when they can be identified, they may be difficult to distinguish from classic hard sweeps (PETER *et al.*, 2012; SCHRIDER *et al.*, 2015). This can be understood from first principles by recognizing that the identification of a standing sweep amounts to recognizing that a particular region of the genome effectively experienced a reduction in effective population size by a factor of *f*, followed by a period of rapid growth back up to the *N*_*e*_ experienced by the bulk of the genome. This task is made difficult by the fact that one has effectively only a single genealogy with which to make this inference, rather imperfect information about its shape and size, and that it shares features both with genealogies expected under a classic hard sweep and under neutrality.

As a result, we suspect that efforts to detect selection from standing variation will continue to be most effective when additional data is available from populations where the allele was not favored or failed to spread for some other reason (INNAN And KIM, 2008; CHEN *et al.*, 2010; ROESTI *et al.*, 2014). If we have good evidence that the allele has spread rapidly, e.g. if the populations are very closely related, then evidence that it is a sweep from standing variation could be gained from demonstrating that the genomic width of the sweep was much smaller than expected and there are too many intermediate frequency haplotypes, given how quickly it would have to have transited through the population. Ancient DNA is also likely to be of value, as we may similarly be able to identify alleles that were at low frequency too recently in the past given observed present day patterns of genetic variation.

## Simulation Details

In order to check our analytical results, we wrote a program to simulate allele frequency trajectories under our model, and then ran either custom written structured coalescent simulations (Figures 3, S1, S2 and S3) or handed these trajectories to *mssel* in order to generate sequence data (Figures 4, 5, 6 and 7).

## Frequency Trajectories

We simulate frequency trajectories under a similar discretized approximation to the diffusion as that used by PRZEWORSKI *et al.* (2005). In order to simulate trajectories conditional on selection having begun when the allele was at frequency *f*, we set *X* (0) = *f*, and simulate allele frequency change forward in time according to

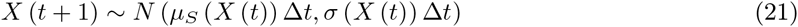

Where

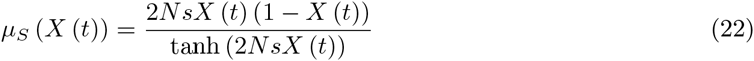

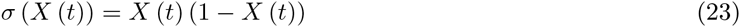

and we take 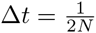so that one time step is equivalent to the duration of one generation in the discrete time Wright-Fisher model, and we have conditioned on the eventual fixation of the allele. To simulate the neutral portion of the frequency trajectory prior to the onset of selection, conditional on the allele having been derived, we take advantage of the time reversibility property of the diffusion process, which dictates that the distribution on the prior history of an allele conditional on being derived and being found at frequency *f* is the same as the future trajectory of an allele that is at frequency *f* and destined to be lost from the population. This allows us to simulate from

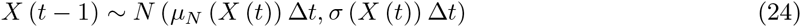

where

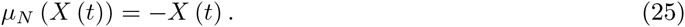

We simply then paste these two trajectories together to give a frequency trajectory that is conditioned on a sweep beginning when the allele is at frequency *f* without any unnatural conditioning on the sweep beginning the *first* time the allele reaches frequency *f*. For simulations intended to check our standing phase calculation independent of the sweep phase, we simply discard the sweep portion of the simulation and retain only the neutral trajectory.

### Genealogy and Recombination Histories

We simulate the genealogy backward in time at the locus of the beneficial allele by allowing a coalescent event to occur between two randomly chosen lineages in generation *t* with probability 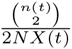, where *n* (*t*) gives the number of lineages existing in generation *t*. Coalescent times obtained from these simulations are then used to generate Figures S1, S2, and S3. We then simulate the history of recombination events which move lineages off of the beneficial background as follows:

We calculate the total time in the genealogy, *T*, and simulate an exp (-*rT*) distance to the first recombination event. A position on the tree (i.e. a branch and a specific generation for the event to occur in) is chosen uniformly at random. This event is accepted with probability 1 − *X* (*t*_*rec*_), where *t*_*rec*_ gives the generation in which the event occurred, otherwise it is ignored. This process is repeated outward away from the focal site until the end of the sequence is reached, and the physical position along the sequence, the branch, and the generation of each recombination event is recorded.

We then generate haplotype identities (i.e. the colorings of Figure 2) as follows. We begin at the root of the tree, and assign each sequence to have the same identity over their entire length. We then move forward in time, from one recombination event to the next, and for each recombination event we assign the chromosomes which subtend it a unique new identity extending from the position where that event occurred out the end of the sequence, overwriting whatever identity previously existed there. We iterate this procedure all the way until the present day, at which point each position on a given chromosome will have an identity specified by the most recent recombination event which falls in between it and the focal site. These haplotype identities are what we use to generate Figure 3.

### Simulated Sequence Data

To simulate sequence data, we simply hand the trajectories simulated as described above to the program mssel (developed by RR Hudson, compiled code is included in supplementary files of this paper). Whereas our simulations of haplotype identity above still represents a somewhat heuristic approximation to the true process, in that it ignores recombination events which do not result in transitions to the alternate background, these simulations of sequence data are exact under the structured coalescent with recombination for our model up the discretization of the diffusion process used for the selected allele. These simulations are used to generate Figures 4, 5, 6 and 7.

#### Code Availability

All custom simulation code was written by JB in R and is provided in Supplementary File 1.

## Appendix

### Incompatibility of Standing Sweep and Classic Strong Selection Models

Consider the small *r* approximation 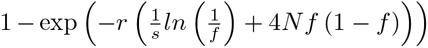 for the recovery of diversity under our standing sweep model, and the approximation

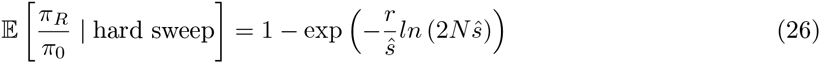

for the hard sweep model. We can set these two expressions equal to one another and solve for ŝ yielding

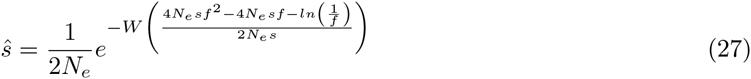

where *W* (*z*) is Lambert ' s W function. This function evaluates to approximately zero for all sensible combinations of parameters under our model, and this fact is responsible for inability to map the effect of sweeps under the standing sweep model to the strong selection hard sweep model.

### Inclusion of New Mutations in Frequency Spectrum

In the main text, we defined *q*(*j* | *k*) as the expected number of segregating mutations present on *j* out of *k* ancestral lineages. Here, we introduce a subscript *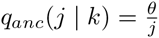* and then define

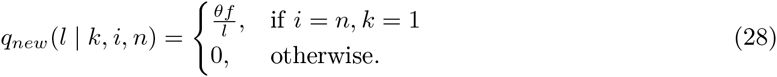

We can then give an improvement upon equation (14) as

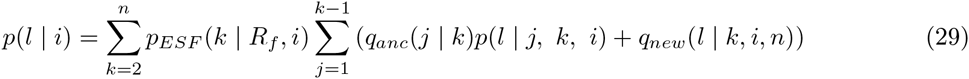

and obtain the normalized version by dividing by their sum.

We can make an improvement upon equation (15) in a similar manner. We first redefine

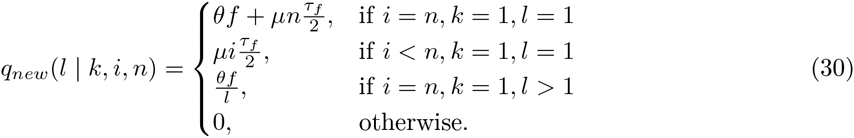

In other words, we allow new mutations to occur during the standing phase provided that there have been no recombination events during this phase, and we also allow new mutations during the sweep phase on all lineages which do not recombine during that phase. Obtaining an improvement over equation (15) is then once again simply a matter of adding the term for new mutations into the previous expression

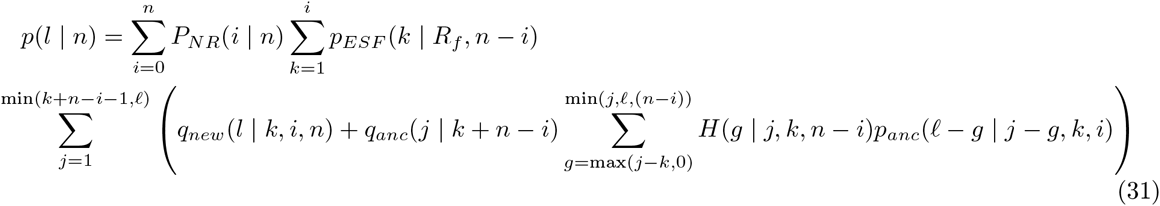

### When Does This Model Apply?

From HERMISSON and PENNINGS (2005), given that one or more adaptive alleles are present in the population and either fixed or destined for fixation *G* generations after an environmental change, and that these alleles were neutral prior to the environmental change, the probability that a population uses material from the standing variation is approximately

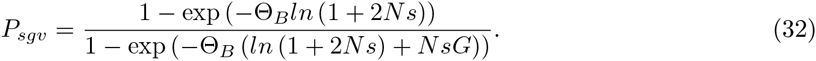

If the population uses material from the standing variation, the probability of finding a single uniquely derived copy of the allele in a sample of *n* lineages is approximately

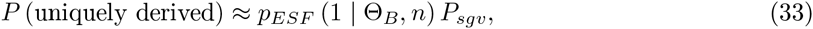

following from the work of (PENNINGS and HERMISSON, 2006a). If this allele was at a frequency less than 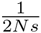 at the moment of the environmental change, the the signature left in polymorphism data will be that of a hard sweep (see PRZEWORSKI *et al.*, 2005). The probability density function for the frequency *f* of a derived allele at the moment of the environmental change, conditional on its eventual fixation, is

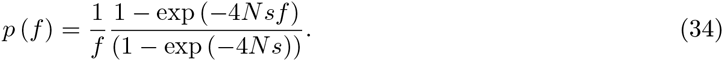

such that we can define the probability that our sweep from standing variation comes from between frequencies *a* and *b* as

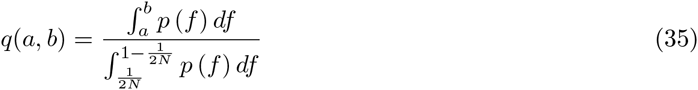

The probability that a sweep of a uniquely derived allele from the standing variation will leave a classic hard sweep signature is therefore

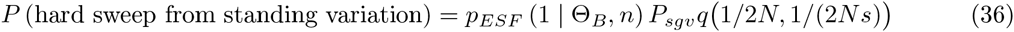

If we take *f* = 0.05 as an approximate upper bound on the frequency from which a uniquely derived sweep from standing variation can be successfully detected, then the probability of such an event is

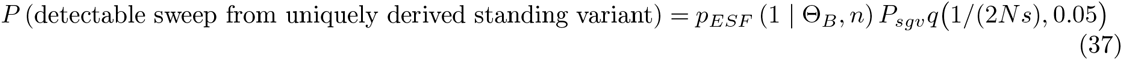

whereas the probability that adaptation proceeds from a uniquely derived standing variant but is essentially undetectable is

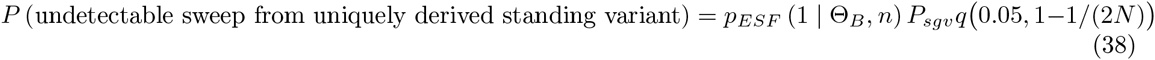

On the other hand, the probability of obtaining a classic hard sweep signature via a new mutation which occurs after the environmental change is

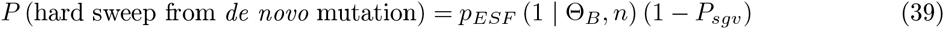

while the probability of a multiple mutation soft sweep, regardless of whether it comes from standing variation or *de novo* mutation is

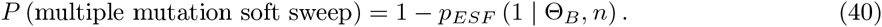

## Acknowledgements

We would like to thank Simon Aeschbacher, Gideon Bradburd, Ivan Juric, Kristin Lee, Alisa Sedghifar, CHENling Xu, and members of the Ross-Ibarra and Schmitt labs at UC Davis for helpful feedback on the work described in this paper.

## 1 Supplementary materials

**Figure S1:**
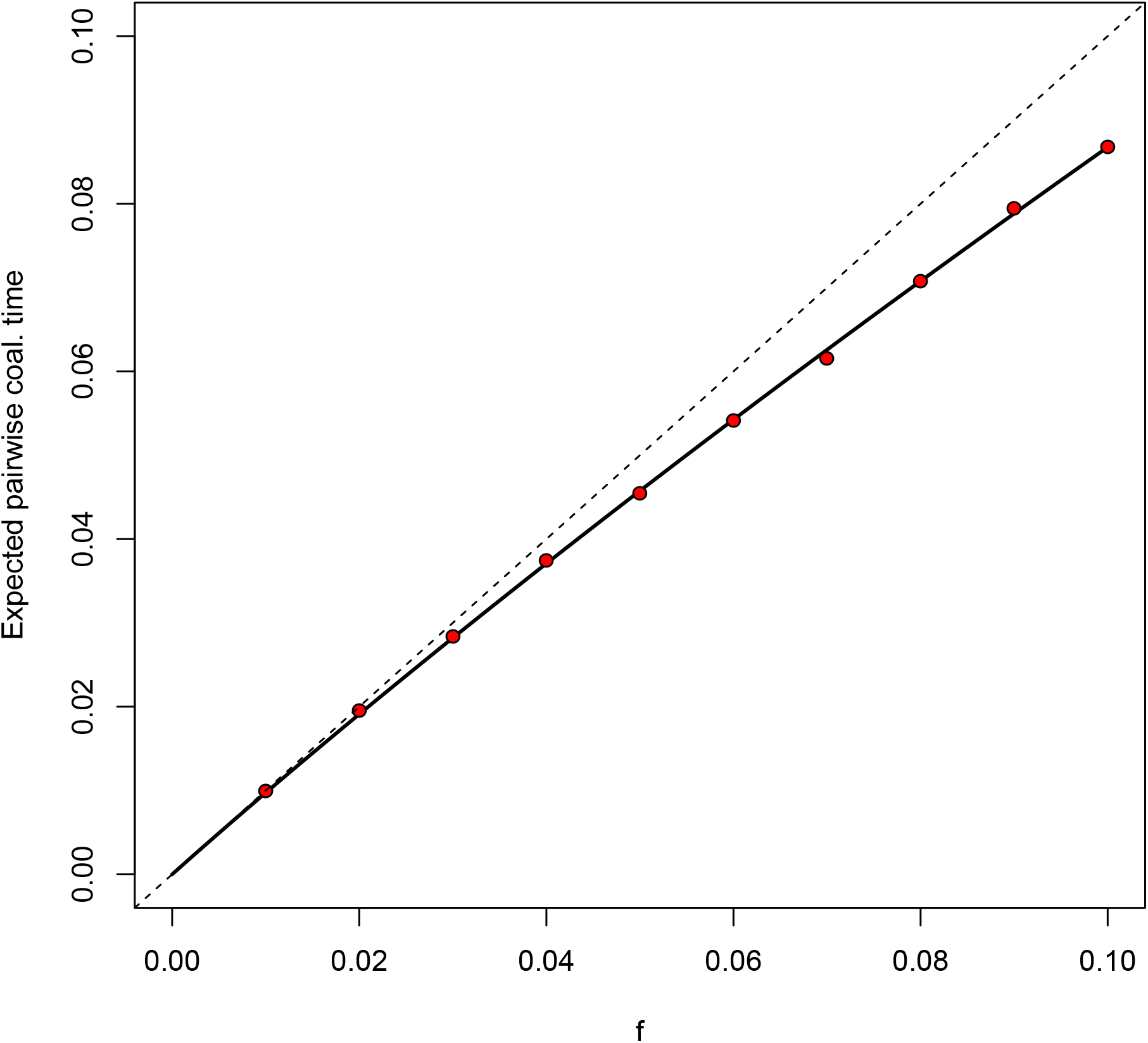
The expected pairwise coalescent time, in units of 2*N* generation, on the background of a derived neutral allele with frequency *f* in the population. The dashed *x* = *y* line shows our approximation 𝔼 [*T*_2_] = 2*N f* The red dots show the mean of our pairwise coalescent simulations featuring an explicit stochastic trajectories. The solid line shows the analytical expectation from GRIFFITHS (2003), equation 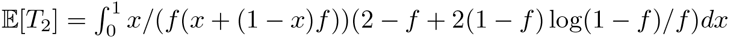.

**Figure S2:**
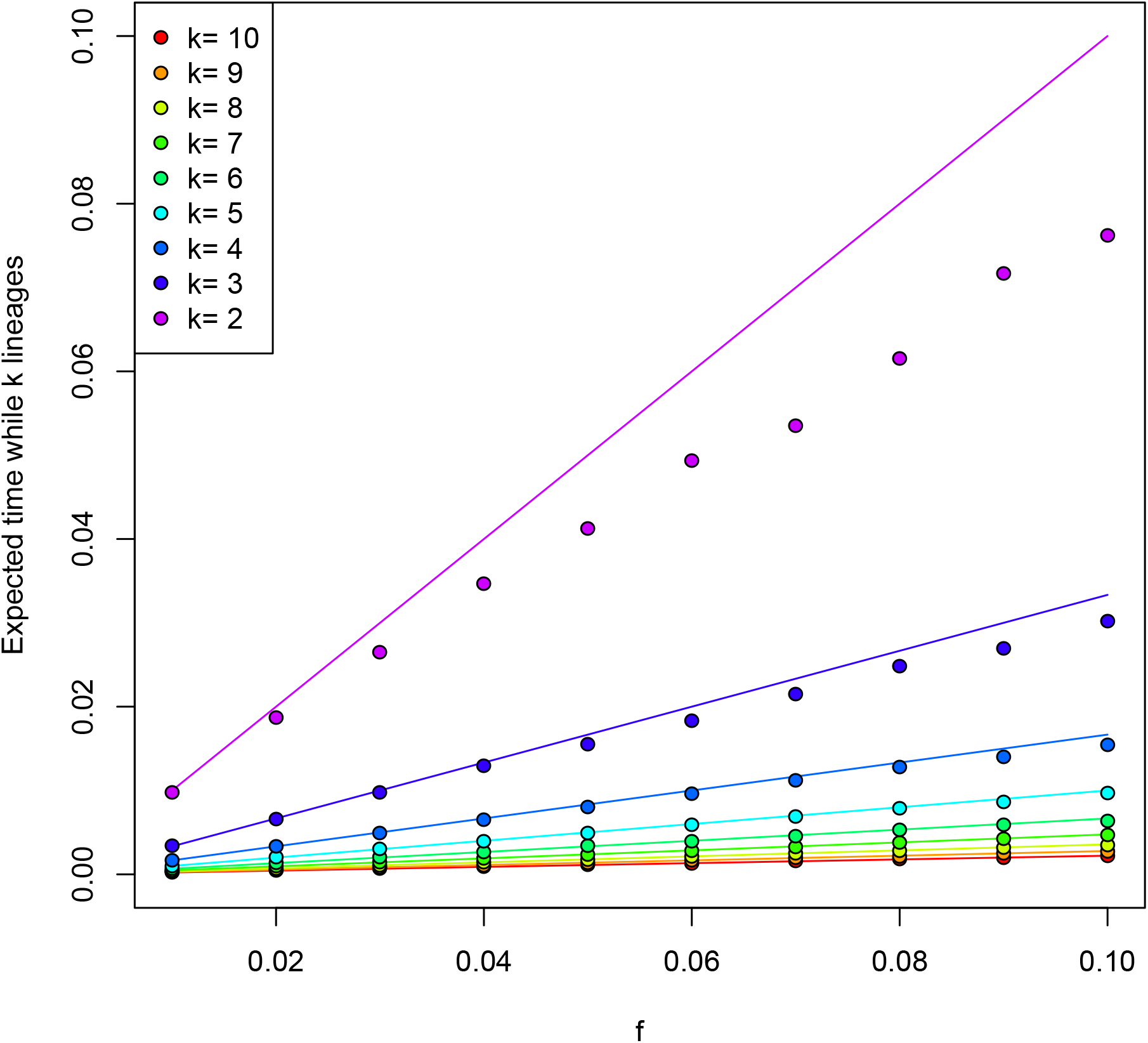
Expected inter-coalescent time intervals for 10 lineages sampled in the current day on the background of a derived neutral allele with frequency *f* in the population. The colored dots give means of *T*_*k*_ from our simulations using stochastic trajectories. The solid lines show the expected coalescent times under our approximation 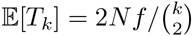 Note the good agreement except for *k* = 2. Presumably our approximation over estimates this time as it fails to acknowledge that the allele is derived, and hence is decreasing in frequency backward in time.

**Figure S3:**
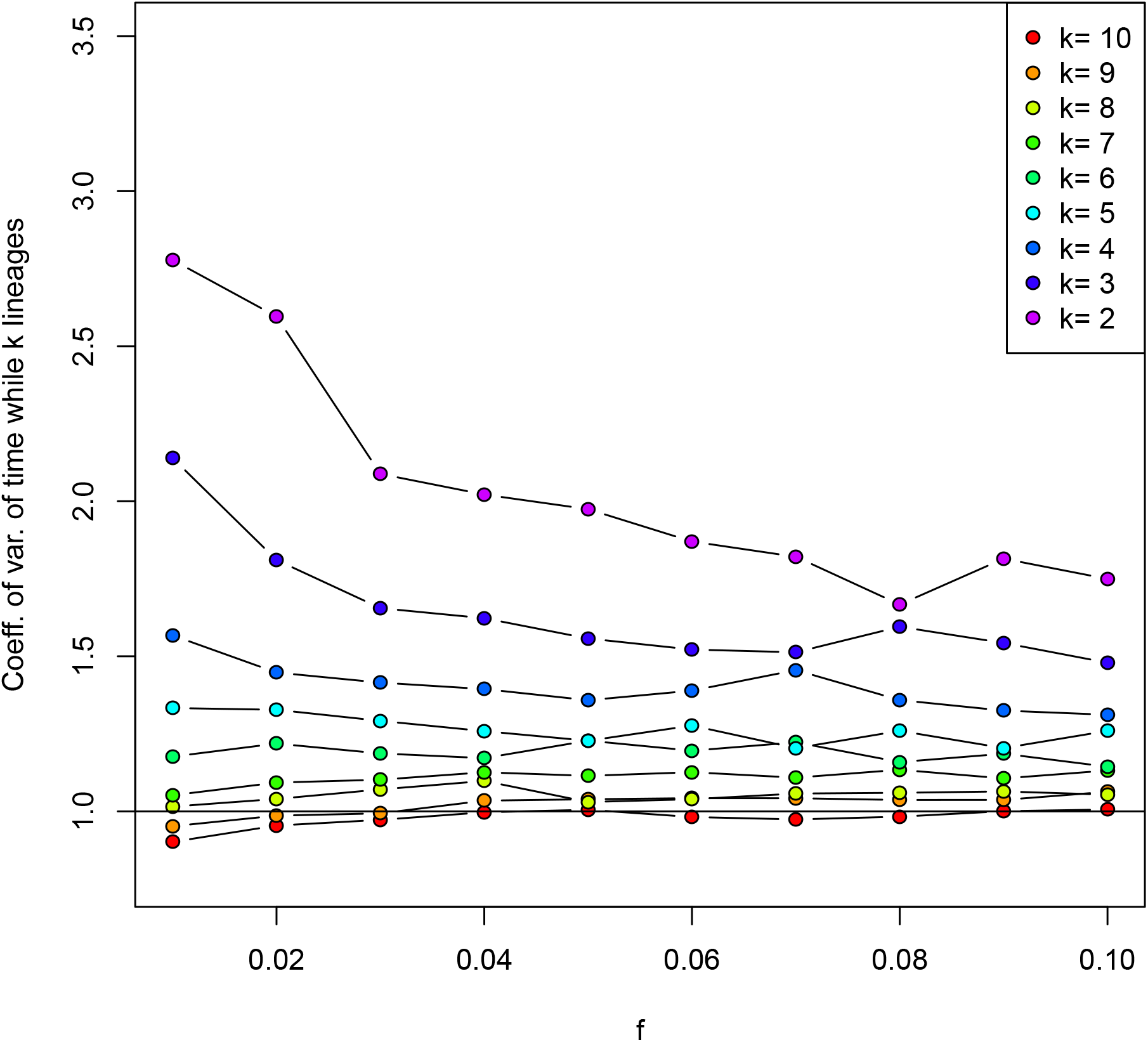
The coefficient of variation (CV) of the inter-coalescent time intervals for 10 lineages sampled in the current day on the background of a derived neutral allele with frequency *f* in the population. The colored dots give the CVs of *T*_*k*_ from our simulations using stochastic trajectories. Under our approximation the coalescent time intervals are exponential, and so have CV=1. The deeper coalescent time-intervals (low *k*s) are more variable than our predictions presumably because neutral trajectories are highly variable so increasing the variance of the coalescent times. More recent time-intervals (high k) are in better agreement with our approximation, as the trajectory with not have strayed far from a frequency *f* a short way back in time.

## References

Andolfatto, P., 2007, December)Hitchhiking effects of recurrent beneficial amino acid substitutions in the Drosophila melanogaster genome. Genome Research 17 (12): 1755–1762.

Bank, C., G. B. Ewing, A. Ferrer-Admettla, M. Foll, and J. D. Jensen, 2014, December)Thinking too positive? Revisitingcurrent methods of populationgenetic selection inference. Trends in Genetics 30 (12): 540–546.

Barrett, R. D. H. and D. Schluter, 2008, January)Adaptation from standing genetic variation. Trends in Ecology & Evolution 23 (1): 38–44.

Barton, N. H., 1998 The effect of hitch-hiking on neutral genealogies. Genetical Research 72 (02).

Chen, H., N. Patterson, and D. Reich, 2010, March)Population differentiation as a test for selective sweeps. Genome Research 20 (3): 393–402.

Colosimo, P. F., 2005, March)Widespread Parallel Evolution in Sticklebacks by Repeated Fixation of Ectodysplasin Alleles. Science (New York, N.Y.) 307 (5717): 1928–1933.

Domingues, V. S., Y.-P. Poh, B. K. Peterson, P. S. Pennings, J. D. Jensen, and H. E. Hoekstra, 2012 Evidence of Adaptation From Ancestral Variation in Young Populations of Beach Mice. Evolution 66 (10): 3209–3223.

Durrett, R. and J. Schweinsberg, 2004, September)Approximating selective sweeps. Theoretical Population Biology 66 (2): 129–138.

Durrett, R. and J. Schweinsberg, 2005, October)A coalescent model for the effect of advantageous mutations on the genealogy of a population. Stochastic Processes and their Applications 115 (10): 1628–1657.

Elyashiv, E., S. Sattath, T. T. Hu, A. Strustovsky, G. McVicker, P. Andolfatto, G. Coop, and G. Sella, 2014 A genomic map of the effects of linked selection in Drosophila. arXiv preprint arXiv:1408.5461.

Etheridge, A., P. Pfaffelhuber, and A. Wakolbinger, 2006, May)An approximate sampling formula under genetic hitchhiking. The Annals of Applied Probability 16 (2): 685–729.

Ewens, W. J., 1972 The Sampling Theory of Selectively Neutral Alleles. Theoretical Population Biology 112: 87–112.

Ewing, G., J. Hermisson, P. Pfaffelhuber, and J. Rudolf, 2011, September)Selective sweeps for recessive alleles and for other modes of dominance. Journal of Mathematical Biology 63 (3): 399–431.

Eyre-Walker, A. and P. D. Keightley, 2009, August)Estimating the Rate of Adaptive Molecular Evolution in the Presence of Slightly Deleterious Mutations and Population Size Change. Molecular Biology and Evolution 26 (9): 2097–2108.

Fay, J. C. and C. I. Wu, 2000, July)Hitchhiking under positive Darwinian selection. Genetics 155 (3): 1405–1413.

Garud, N. R., P. W. Messer, E. O. Buzbas, and D. a. Petrov, 2015, February)Recent Selective Sweeps in North American Drosophila melanogaster Show Signatures of Soft Sweeps. PLoS Genetics 11 (2): e1005004.

Garud, N. R. and N. A. Rosenberg, 2015, April)Theoretical Population Biology. Theoretical population biology: 1–8.

Griffiths, R. C., 1980, February)Lines of descent in the diffusion approximation of neutral Wright-Fisher models. Theoretical population biology 17 (1): 37–50.

Griffiths, R. C., 2003, September)The frequency spectrum of a mutation, and its age, in a general diffusion model. Theoretical Population Biology 64 (2): 241–251.

Griffiths, R. C. and S. Tavare, 1998, January)The age of a mutation in a general coalescent tree. Stochastic Models 14 (1-2): 273–295.

Griffiths, R. C. and S. Tavare, 1999 The ages of mutations in gene trees. The Annals of Applied Probability: 567–590.

Hermisson, J. and P. S. Pennings, 2005 Soft sweeps: molecular population genetics of adaptation from standing genetic variation. Genetics 169 (4): 2335–2352.

Hermisson, J. and P. Pfaffelhuber, 2008 The pattern of genetic hitchhiking under recurrent mutation. Electronic Journal of Probability 13: 2069–2106.

Innan, H. and Y. Kim, 2004, July)Pattern of polymorphism after strong artificial selection in a domestication event. Proceedings of the National Academy of Sciences 101 (29): 10667–10672.

Innan, H. and Y. Kim, 2008, June)Detecting Local Adaptation Using the Joint Sampling of Polymorphism Data in the Parental and Derived Populations. Genetics 179 (3): 1713–1720.

Innan, H. and M. Nordborg, 2003, September)The extent of linkage disequilibrium and haplotype sharing around a polymorphic site. Genetics 165 (1): 437–444.

Jensen, J. D., 2014 On the unfounded enthusiasm for soft selective sweeps. Nature Communications 5: 5281.

Jones, B. L., T. O. Raga, A. Liebert, P. Zmarz, E. Bekele, E. T. Danielsen, A. K. Olsen, N. Bradman, J. T. Troelsen, and D. M. Swallow, 2013, September)REPOR TDiversity of Lactase Persistence Alleles in Ethiopia: Signature of a Soft Selective Sweep. American journal of human genetics 93 (3): 538–544.

Kaplan, N. L., R. R. Hudson, and C. H. Langley, 1989 The “hitchhiking effect” revisited. Genetics 123 (4): 887–899.

Karasov, T., P. W. Messer, and D. a. Petrov, 2010, June)Evidence that Adaptation in Drosophila Is Not Limited by Mutation at Single Sites. PLoS Genetics 6 (6): e1000924.

Kimura, M. and J. F. Crow, 1964, April)THE NUMBER OF ALLELES THAT CAN BE MAINTAINED IN A FINITE POPULATION. Genetics 49: 725–738.

Li, H. and R. Durbin, 2011, July)Inference of human population history from individual whole-genome sequences. Nature 475 (7357): 493–496.

Magwire, M. M., F. Bayer, C. L. Webster, C. Cao, and F. M. Jiggins, 2011, October)Successive Increases in the Resistance of Drosophila to Viral Infection through a Transposon Insertion Followed by a Duplication. PLoS Genetics 7 (10): e1002337.

Maynard Smith J., and J. Haigh, 1974, February)The hitch-hiking effect of a favourable gene. Genetics Research 23 (1): 23–35.

McVean, G., 2006, December)The Structure of Linkage Disequilibrium Around a Selective Sweep. Genetics 175 (3): 1395–1406.

Menozzi, P., M. A. Shi, A. Lougarre, Z. H. Tang, and D. Fournier, 2004, February)Mutations of acetylcholinesterase which confer insecticide resistance in Drosophila melanogaster populations. BMC evolutionary biology 4: 4.

Messer, P. W. and R. A. Neher, 2012, June)Estimating the strength of selective sweeps from deep population diversity data. Genetics 191 (2): 593–605.

Messer, P. W. and D. a. Petrov, 2013, November)Population genomics of rapid adaptation by soft selective sweeps. Trends in Ecology & Evolution 28 (11): 659–669.

Nair, S., D. Nash, D. Sudimack, A. Jaidee, M. Barends, A.-C. Uhlemann, S. Krishna, F. c. c. o. Nosten, and T. J. C. Anderson, 2006, November)Recurrent gene amplification and soft selective sweeps during evolution of multidrug resistance in malaria parasites. Molecular Biology and Evolution 24 (2): 562–573.

Nielsen, R., S. Williamson, Y. Kim, M. J. Hubisz, A. G. Clark, and C. Bustamante, 2005, November)Genomic scans for selective sweeps using SNP data. Genome Research 15 (11): 1566–1575.

Patterson, N. J., 2005 How old is the most recent ancestor of two copies of an allele? Genetics 169 (2): 1093–1104.

Paul, J. S., M. Steinrücken, and Y. S. Song, 2011, April)An accurate sequentially Markov conditional sampling distribution for the coalescent with recombination. Genetics 187 (4): 1115–1128.

Pennings, P. S. and J. Hermisson, 2006a Soft sweeps II–molecular population genetics of adaptation from recurrent mutation or migration. Molecular Biology and Evolution 23 (5): 1076–1084.

Pennings, P. S. and J. Hermisson, 2006b Soft sweeps III: the signature of positive selection from recurrent mutation. PLoS Genetics 2 (12): e186.

Peter, B. M., E. Huerta-Sanchez, and R. Nielsen, 2012, October)Distinguishing between Selective Sweeps from Standing Variation and from a De Novo Mutation. PLoS Genetics 8 (10): e1003011.

Pokalyuk, C., 2012, January)The effect of recurrent mutation on the linkage disequilibrium under a selective sweep. Journal of Mathematical Biology 64 (1-2): 291–317.

Przeworski, M., G. Coop, and J. D. Wall, 2005 The signature of positive selection on standing genetic variation. Evolution; international journal of organic evolution 59 (11): 2312–2323.

Ralph, P. and G. Coop, 2010 Parallel adaptation: one or many waves of advance of an advantageous allele? Genetics 186 (2): 647–668.

Rannala, B., 1997, December)On the genealogy of a rare allele. Theoretical population biology 52 (3): 216–223.

Rasmussen, M. D., M. J. Hubisz, I. Gronau, and A. Siepel, 2014, May)Genome-Wide Inference of Ancestral Recombination Graphs. PLoS Genetics 10 (5): e1004342.

Roesti, M., S. Gavrilets, A. P. Hendry, W. Salzburger, and D. Berner, 2014, August)The genomic signature of parallel adaptation from shared genetic variation. Molecular Ecology 23 (16): 3944–3956.

Sabeti, P. C., D. E. Reich, J. M. Higgins, H. Z. P. Levine, D. J. Richter, S. F. Schaffner, S. B. Gabriel, J. V. Platko, N. J. Patterson, G. J. McDonald, H. C. Ackerman, S. J. Campbell, D. Altshuler, R. Cooper, D. Kwiatkowski, R. Ward, and E. S. Lander, 2002, October)Detecting recent positive selection in the human genome from haplotype structure. Nature 419 (6909): 832–837.

Salgueiro, P., J. L. Vicente, C. Ferreira, V. Teófilo, A. Galvão, V. E. do Rosário, P. Cravo, and J. Pinto, 2010 Tracing the origins and signatures of selection of antifolate resistance in island populations of Plasmodium falciparum. BMC Infectious Diseases 10 (1): 163.

Schmidt, J. M., R. T. Good, B. Appleton, J. Sherrard, G. C. Raymant, M. R. Bogwitz, J. Martin, P. J. Daborn, M. E. Goddard, P. Batterham, and C. Robin, 2010, June)Copy Number Variation and Transposable Elements Feature in Recent, Ongoing Adaptation at the Cyp6g1 Locus. PLoS Genetics 6 (6): e1000998.

Schrider, D. R., F. K. Mendes, M. W. Hahn, and A. D. Kern, 2015, May)Soft Shoulders Ahead: Spurious Signatures of Soft and Partial Selective Sweeps Result from Linked Hard Sweeps. Genetics 200 (1): 267–284.

Schweinsberg, J. and R. Durrett, 2005, August)Random partitions approximating the coalescence of lineages during a selective sweep. The Annals of Applied Probability 15 (3): 1591–1651.

Sheehan, S., K. Harris, and Y. S. Song, 2013, April)Estimating Variable Effective Population Sizes From Multiple Genomes: A Sequentially Markov Conditional Sampling Distribution Approach. Genetics.

Studer, A., Q. Zhao, J. Ross-Ibarra, and J. Doebley, 2011, November)Identification of a functional transposon insertion in the maize domestication gene tb1. Nature Genetics 43 (11): 1160–1163.

Voight, B. F., S. Kudaravalli, X. Wen, and J. K. Pritchard, 2006 A Map of Recent Positive Selection in the Human Genome. PLoS Biology 4 (3): e72.

Watterson, G. A., 1975 On the number of segregating sites in genetical models without recombination. Theoretical population biology 7 (2): 256–276.

Watterson, G. A., 1984 Lines of descent and the coalescent. Theoretical population biology 26 (1): 77–92.

Wiehe, T. H. and W. Stephan, 1993, July)Analysis of a genetic hitchhiking model, and its application to DNA polymorphism data from Drosophila melanogaster. Molecular Biology and Evolution 10 (4): 842–854.

Wilson, B. A., D. a. Petrov, and P. W. Messer, 2014, October)Soft selective sweeps in complex demographic scenarios. Genetics 198 (2): 669–684.

Wiuf, C., 2000, August)On the genealogy of a sample of neutral rare alleles. Theoretical Population Biology 58 (1): 61–75.

Wiuf, C. and P. Donnelly, 1999 Conditional genealogies and the age of a neutral mutant. Theoretical Population Biology 56 (2): 183–201.

